# GenePT: A Simple But Effective Foundation Model for Genes and Cells Built From ChatGPT

**DOI:** 10.1101/2023.10.16.562533

**Authors:** Yiqun Chen, James Zou

## Abstract

There has been significant recent progress in leveraging large-scale gene expression data to develop foundation models for single-cell biology. Models such as Geneformer and scGPT implicitly learn gene and cellular functions from the gene expression profiles of millions of cells, which requires extensive data curation and resource-intensive training. Here we explore a much simpler alternative by leveraging ChatGPT embeddings of genes based on literature. Our proposal, GenePT, uses NCBI text descriptions of individual genes with GPT-3.5 to generate gene embeddings. From there, GenePT generates single-cell embeddings in two ways: (i) by averaging the gene embeddings, weighted by each gene’s expression level; or (ii) by creating a sentence embedding for each cell, using gene names ordered by the expression level. Without the need for dataset curation and additional pretraining, GenePT is efficient and easy to use. On many downstream tasks used to evaluate recent single-cell foundation models — e.g., classifying gene properties and cell types — GenePT achieves comparable, and often better, performance than Geneformer and other models. GenePT demonstrates that large language model embedding of literature is a simple and effective path for biological foundation models.

## 1 Introduction

Recently, the field of single-cell biology has seen a surge in interest and efforts to develop foundation models, i.e., models designed to learn embeddings of genes and cells to facilitate various downstream analyses. Several methods, such as Geneformer [1] and scGPT [2], have been recently proposed to tackle this challenge. At a conceptual level, they adopt similar recipes that consist of the following steps:

1. Adopt a deep learning architecture (often from the transformer family [3]).
2. Gather extensive single-cell gene expression datasets for pretraining the model in a self-supervised manner (e.g., by imputing some masked-out expression values). The trained encoder maps input genes and cells to a high-dimensional embedding vector encapsulating the underlying biology.
3. For downstream tasks, one can optionally utilize a modest amount of task-specific data to fine-tune the model, boosting its predictive capabilities.

Notably, the approach outlined above derives embeddings *only* from gene expression datasets, without making any use of the literature and pre-existing knowledge about a gene. While this strategy has shown some success in applications to single-cell transcriptomics data and tasks, it has several limitations. First, the computational power and time required to collect and process large-scale single-cell transcriptomics data used for pretraining (Step 2 above) can be prohibitive, particularly when researchers desire early signal detection and rapid iterations. Furthermore, the signals from extracted embeddings are heavily dependent on the gene expression data used in Step 2, which doesn’t take advantage of the vast research and literature summarizing the functionalities of a gene, potentially leading to sample inefficiency and suboptimal results in certain applications. Therefore, in this study, we explored an alternative, complementary approach and investigated the feasibility of encoding the biology of genes and cells using natural language.

The intuition for our approach is as follows: large-language models (LLMs) such as GPT-3.5 and GPT-4 have been trained on extensive text corpus [4], including biomedical literature, and have demonstrated remarkable ability in understanding, reasoning, and even generating biomedical text [5–8]. Consequently, we hypothesize that LLM-derived embeddings of gene summaries and functionalities — which often are curated from a broad spectrum of experiments and studies — might more directly capture the underlying biology.

We introduced GenePT — a method that represents genes and cells by utilizing OpenAI’s ChatGPT text embedding API services [9]. We evaluated the generated embeddings on several biologically driven tasks and our findings reveal that GenePT exhibits performance comparable to, and sometimes surpassing, specially designed models such as Geneformer across a diverse set of downstream tasks. GenePT offers several advantages to single-cell RNA-seq-based foundation models: (i) it performs better on several biological tasks; (ii) it doesn’t require expensive single-cell curation and additional pretraining; and (iii) it’s very simple to use and to generate gene and cell embeddings. GenePT uses LLM-based embeddings which is an orthogonal source of information compared to the expression-based representations; this suggests a promising new direction of combining these two approaches.

## 2 Related Work

### 2.1 Foundation models for single-cell transcriptomics

Foundation models have shown unprecedented performance for a myriad of tasks including text classification, question answering, and text generation [10]. Efforts have naturally been made to adapt these models to tackle tasks in biology, especially in the field of single-cell transcriptomics [1, 2, 11]. Examples of such efforts include cell type annotation, where a cell is labelled based on its biological identity [12, 13]); gene functional and regulatory network inference, where the functionality of individual genes and clustered gene groups are examined [2, 14]; and sample integration [13], which accounts for transcript abundance influenced primarily by technical noise instead of underlying biology.

With the advent of large-scale, open-source expression datasets such as Gene Expression Omnibus [15] and the Human Cell Atlas [16], several models have been trained on such data. The aspiration behind these models is to craft a foundational model for single-cell transcriptomics, analogous to foundational models in natural language processing. These models are intended to display broad capabilities across an array of biological tasks rather than just a niche subset. For instance, Geneformer [1] employs extensive pretraining on the ranks of gene expression levels through masked token prediction across 30 million cells collected from a wide range of sources using a transformer architecture. It shows good performance in tasks ranging from understanding network dynamics to deciphering network hierarchy. Another noteworthy model is scGPT [2]: it hinges on generative pretraining (with gene expression prediction as the task) and used 33 million cells from the CELLxGENE collection for training [17]. Its capabilities are demonstrated through downstream evaluations in perturbation prediction, batch integration, and cell type annotation.

### 2.2 Using language models for cell biology

Pioneering work in applying language models to gene and cell biology aims to represent the semantics of biomedical terms by training co-occurrence-based neural network embeddings that map individual terms (e.g., gene names) to vectors [18–20]. Recently, researchers have begun exploring the use of LLMs for biomedically-focused tasks, leveraging their capability to encode information from the entire input text. This approach allows for more nuanced and dynamic representations. For example, Hou and Ji [21] employed ChatGPT for cell type annotation; Wysocki et al. [22] investigated biomedical meanings encoded by BioBERT and BioMegatron embeddings; and Ye et al. [23] utilized instruction fine-tuning to achieve competitive results on graph data task benchmarks with an LLM. Compared to prior works that directly query LLMs for biological tasks, our method solely utilizes the input descriptions of each gene (which can be sourced from high-quality databases such as NCBI [24]) and the embedding model of LLMs, which suffers less from problems such as hallucination. While our paper is under preparation, Levine et al. [25] has independently embarked on a conceptually related approach to ours, where each cell is transformed into a sequence of gene names, ranked by expression level and truncated at top 100 genes. The emphasis of their paper, however, is on generating new cells conditional on cell types.

### 2.3 Deciphering natural language embeddings

Understanding how large-scale unsupervised representations capture linguistic nuances is a central research question in Natural Language Processing (NLP). One avenue of exploration, sometimes referred to as “probes”, trains supervised models to predict downstream properties from the language model embeddings [26–28]. These techniques have achieved impressive accuracy across NLP tasks, suggesting that the embeddings exhibit a substantial amount of understanding of input attributes.

The success of probing and foundational models in biology inspires our primary research questions (RQs):

RQ1: Do natural language embeddings of genes capture the intrinsic biological functioalities of a gene?

RQ2: Do natural language embeddings of cells capture the underlying biology of a cell?

In addressing these RQs, our study makes the following contribution to the literature: we show that natural language embeddings of gene functions — such as summaries readily available from sources like the NCBI gene database [29] — successfully encapsulate the underlying biological relationships and insights associated with genes, when assessed on biologically relevant tasks. Moreover, for single cells, language model embeddings of the gene names, ordered by expression levels, encode substantial biological signals that can be used, e.g., for cell type annotation.

## 3 Methods

### 3.1 Data Collection and Transformation

To obtain embeddings for genes most pertinent to single-cell transcriptomics studies, we began with unifying the list of gene names provided in Geneformer [1] and scGPT [2]. The selection of these genes was informed by their expression levels across the pretraining datasets. In Geneformer cases, the genes were represented as Ensembl IDs rather than gene names, and we used the mygene package [30] for conversion, retaining in successful look-up of more than 90% of the Ensembl IDs. Additionally, we incorporated genes detected in our downstream application datasets, totalling around 33,000 genes. For each gene, we extracted its information from the NCBI gene database’s summary section after removing hyperlinks and date information. GPT-3.5 (text-embedding-ada-002) embeddings were obtained for the summaries for each gene (mean: 73 words; interquartile range: 25–116). Each embedding has a dimension of 1,536, which serves as gene representations (see Figure 1). Moreover, we mapped around 60,000 additional gene name aliases to the NCBI summary embedding using the HGNC database [31]. We conducted sensitivity analyses using three different levels of content input for gene summaries: gene names only, gene names and gene summaries, and all summary card information (see details in Appendix A).

**Fig. 1:**
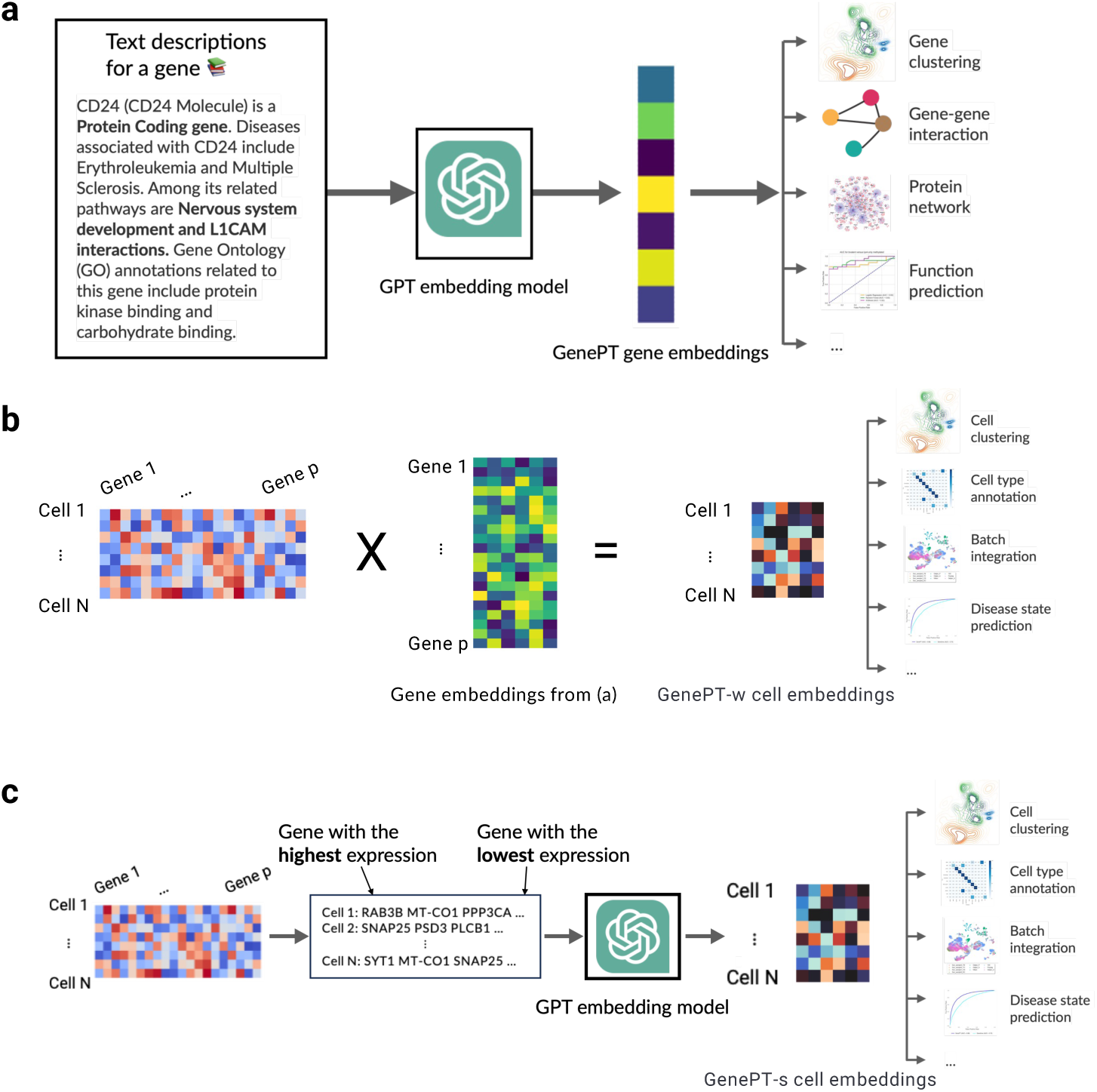
An overview of the GenePT framework. **(a)** For each gene, we extract its corresponding text summary from NCBI and use GPT-3.5 text embedding as its representation. **(b)** In the GenePT-w cell embeddings framework, we average gene embeddings from step (a), weighted by their cell expression levels, and normalize these cell embeddings to a unit *ℓ*_2_ norm. **(c)** In the GenePT-s cell embeddings framework, each cell from the input single-cell data is translated into a natural language sentence based on ranked gene expressions, and the GPT-3.5 embedding of the entire sentence is used to represent the cell.

In addition to embedding the gene summaries using GPT-3.5, we conducted comparisons with alternative embedding methods, such as gene summary embeddings using the open-source biomedical language models such as BioLinkBert [32] and gene-expression-derived embeddings such as Gene2vec [18] and Geneformer [1].

To encode information at the cellular level, we developed two distinct approaches: GenePT-w (w for weighted) and GenePT-s (s for sentence). In both approaches, we first normalize and transform the scRNA-seq data as implemented in the scanpy package as follows: firstly, we row-normalize the count matrix so that each cell has 10,000 observed RNA transcripts, followed by a log(1 + *x*) transformation of each matrix entry.

To construct GenePT-w embeddings, we first take a weighted average of the GenePT gene embeddings, where the weight is determined by the normalized expressions of each gene, and then normalize the embedding to have unit *ℓ*_2_ norm (see Figure 1(b)). This approach leverages the rich context of each gene embedding, but is limited by the simplicity of the weighted average. Alternatively, we can represent cells using natural language sentences by creating a sequence of gene names analogous to sentences. Here, the sequence is ordered by descending normalized expression levels, omitting genes with zero counts. We then pass this sentence representation for each cell to GPT-3.5 to obtain GenePT-s embeddings (see Figure 1(c)).

### 3.2 Downstream gene-level and cell-level applications

Geneformer and scGPT demonstrate the biological value offered by their foundation models using several downstream gene-level and cell-level tasks. In this paper, we evaluated the performance of GenePT on the same downstream applications wherever possible to compare GenePT with Geneformer and other single-cell foundation models. In particular, for gene-level tasks, we primarily contrast our results with those from Geneformer, Gene2vec, and scGPT. This is because their results on the same datasets have been previously documented without the need to re-train or fine-tune. Regarding cell-level tasks, we leveraged *pretrained* embeddings from both Geneformer and scGPT.

The details for the gene-level and cell-level applications are as follows:

- Gene-level Tasks:

– Gene Functionality Class Prediction: This is a multi-class prediction challenge based on the 15 most common functional gene classes. Labels for these classes were curated as part of the Geneformer paper.
– Gene Property Prediction Task: This involves four binary classification tasks using open-source data provided in Theodoris et al. [1]: Distinguishing previously identified dosage-sensitive from dosage-insensitive transcription factors. Differentiating between bivalent and non-methylated genes. Differentiating between Lys4-only-methylated and non-methylated genes. Distinguishing long-range from short-range transcription factors (TFs).
– Gene-Gene Interaction Prediction: We utilized a benchmark for gene-gene interaction (GGI) based on shared gene ontology annotations published by Du et al. [18]. The training and test datasets include over 200,000 pairs of examples in the tuple (gene 1, gene 2, label), where the binary label indicates whether a pair of genes is known to interact.
– Protein-Protein Interaction Prediction: We assessed the ability to predict protein-protein interactions (PPI) using GenePT embeddings with the following three datasets: (i) The human binary protein interactions (HuRI) dataset collected by Luck et al. [33] through screening with multiple PPI assays; (ii) comprehensive binary protein-protein interactions (Lit-BM) that are supported by at least two traceable pieces of evidence [34]; and (iii) tissue-specific protein-protein functional interaction networks derived by Greene et al. [35]. These PPI datasets contain tuples in the form of (protein 1, protein 2, binary label). The binary label indicates whether there’s an observed interaction between the two proteins. We first converted the proteome identifiers for proteins into gene names using the UniProt conversion tool [36]. If multiple genes were returned, we randomly selected one. Since only positive interactions were reported in the HuRI and Lit-BM dataset, we constructed an equal amount of negative data by randomly sampling pairs of proteins examined in Luck et al. [33] that weren’t reported as interacting pairs. We explored the potential utility of creating context-dependent embeddings by providing context-dependent gene descriptions for protein-protein interaction tasks in Appendix B.5.
– Unsupervised Exploration of Gene Programs: To examine the interaction between genes, we constructed a similarity network of gene-gene interactions using GenePT embeddings from a dataset of human immune tissues [37]. Our validation process follows that of Cui et al. [2] and consists of the following steps: 1. constructing gene networks based on the cosine similarities among the highly variable genes; 2. applying unsupervised Louvain clustering [38] to derive gene programs; and 3. qualitatively comparing the trends of highlighted gene programs with their cell-specific expression levels.
- Cell-level tasks:

– Assessing Association Between Embeddings and the Underlying Cell States: Here, we considered the following test datasets representing cells from circulatory systems (Aorta, a random 20% subset of data originally published in Li et al. [39]) comprising 11 cell types; and Artery [40] with 10 cell types), bone tissues (Bones [41] with 7 cell types; Myeloid [42] containing 3 annotated cancer types
and 11 cell types across 13,468 cells)), the Pancreas [37] (containing 11 annotated cell types across 4,218 cells), and immune cells collected from healthy individuals and patients with Multiple Sclerosis [43], totalling 18 annotated cell types and 12 donors across 3,430 cells. For each dataset and its associated metadata annotation, we applied *k*-means clustering on the *pretrained* GenePT, Geneformer, or the scGPT embeddings to obtain clusters matching the classes in the metadata annotations. We select the number of clusters *k* to match the number of classes in the metadata annotation. We then computed the Adjusted Rand Index (ARI) and Adjusted Mutual Information (AMI) to evaluate the concordance between derived cluster labels and the true metadata labels. A higher alignment, indicated by higher values of ARI or AMI, between the inferred and actual labels, suggests that the embedding captures more biological structure and signals. We also calculated the Average Silhouette Width (ASW) using the true annotations of original samples to assess the cohesion and separation of the clusters.
– Context Awareness and Batch Integration: Pretrained single-cell foundation models have been demonstrated to be robust against common batch-dependent technical artifacts while still encoding the underlying biological context. We assessed whether GenePT-s embeddings were impacted by common batch effects such as patient variability on two datasets used in Theodoris et al. [1]: the cardiomyocyte dataset originally published by Chaffin et al. [44], and the Aorta dataset originally published in Li et al. [39].

## 4 Results

### 4.1 GenePT embeddings capture underlying gene functionality

In Figure 2(a), we display a 2D UMAP of the GenePT embeddings (using the text-embedding-ada-002 model), for over 34,000 genes that belong to the top 15 most prevalent functional classes (see Table B2 in Appendix B for detailed class breakdown). The UMAP reveals distinct clusters when coloured by various gene functionality groups, implying that GenePT embeddings encode the functions of the genes. This confirms that language model embedding retains key biological information, as functionality is frequently found in NCBI gene summaries. To evaluate the observations in Figure 2(a) more quantitatively, we further divided the genes into a 70%/30% train/test split and evaluated the prediction accuracy using an *ℓ*_2_ regularized logistic regression on the 15 classes. The predicted functional class aligns with the true annotation well, with an overall accuracy of 96% and high class-specific accuracies and only minor misclassifications between closely related functional groups like lincRNA, lncRNA, and processed transcripts (see Figure 2(b)).

**Fig. 2:**
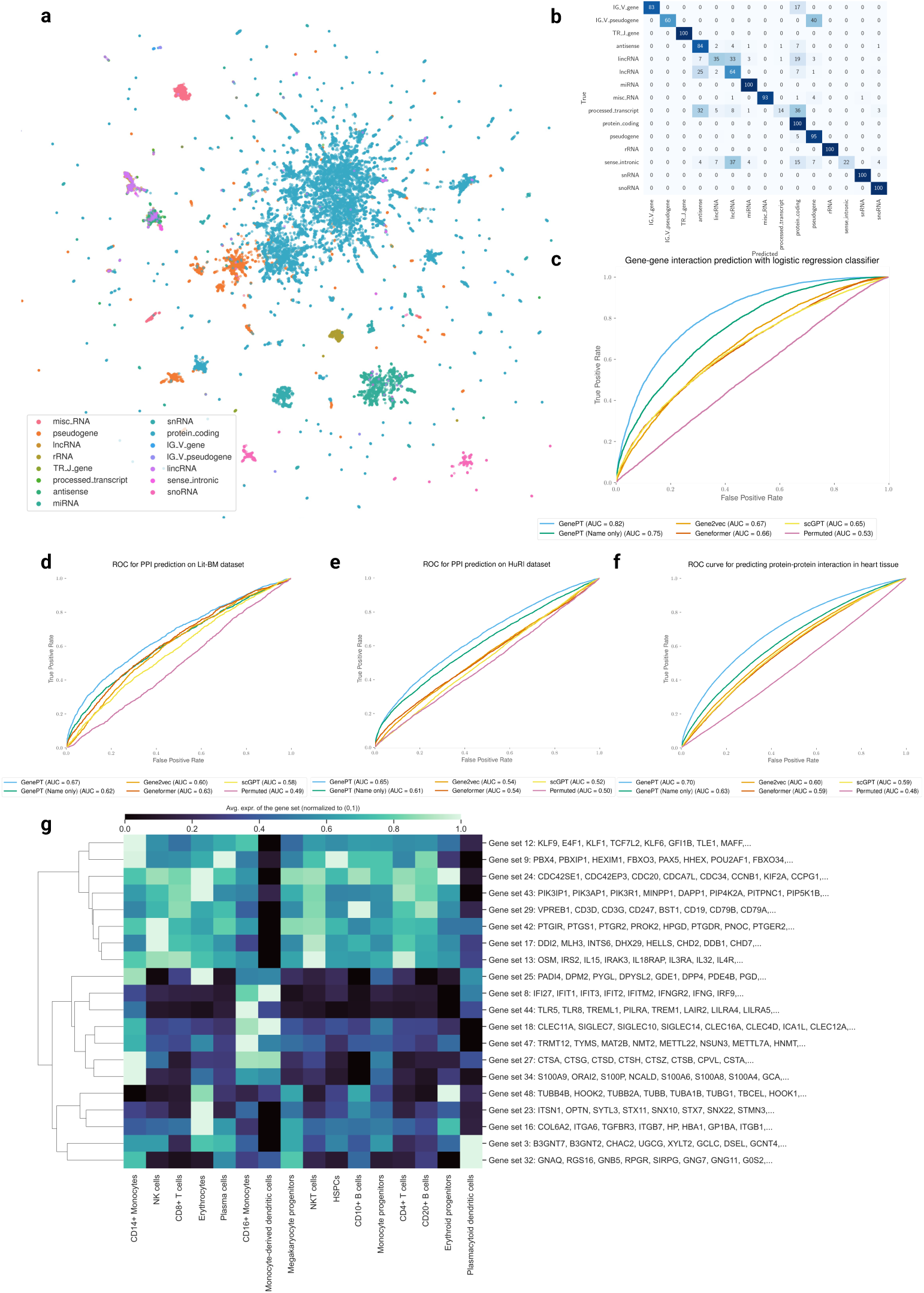
GenePT gene embeddings encode underlying biology. **(a)** 2D UMAP visualization of GenePT embeddings, colored by different gene functionality groups. **(b)** Confusion matrix of gene function prediction utilizing GenePT embeddings, combined with an *ℓ*_2_-regularized logistic regression on a randomly held-out 30% test set. **(c)** Prediction accuracy on a gene-gene interaction benchmark dataset derived from GEO expression data [18]. **(d)** Prediction accuracy for protein-protein interactions verified by high-quality binary literature datasets [34]. **(e)** Prediction accuracy on the human binary protein interactions dataset [33]. **(f)** Prediction accuracy on human heart tissue protein-protein functional interactions [35]. **(g)** Cell-type specific activation among GenePT-embeddings-extracted gene programs (a random subset of genes is displayed for each program) in a human immune tissue dataset [37]. The patterns of average gene expressions for identified gene programs in different cell types are congruent with those previously identified in Cui et al. [2].

We further assessed the efficacy of GenePT embeddings in predicting gene-gene interactions (GGI) in Figure 2(c). We compared the ROC-AUC for three methods on the test GGI dataset provided in Du et al. [18], derived from shared Gene Ontology (GO) annotations: (i) sum of the GenePT embedding of two genes with an *ℓ*_2_-regularized logistic classifier (LR), yielding an AUC of 0.82; (ii) sum of the Gene2Vec/scGPT/Geneformer pretrained embeddings with an LR classifier (resulting in AUC of 0.65–0.67); and (iii) sum of two random embeddings (*d* = 1, 536, same dimension as GenePT) paired with an LR classifier, which served as a negative control (an AUC of 0.51). As shown in Figure 2(c), GenePT embeddings considerably enhance performance when compared to the other single-cell foundation models derived embeddings using the same downstream classifier. Even when leveraging a more intricate deep neural network, Du et al. [18] reported an AUC of 0.77, underscoring the competitive edge of GenePT in this task.

Next, we evaluated the ability to predict protein-protein interactions (PPI) using GenePT gene embeddings, as depicted in Figure 2(d)–(f). We compared the ROC-AUC (refer to Appendix B.3 results for the precision-recall curve) for three methods across three distinct PPI datasets: those derived from the literature (panel (d)), comprehensive assays (panel (e)), and biophysical contact annotations (panel (f)). For all three datasets, using the sum of the GenePT embeddings of two genes as input, combined with an *ℓ*_2_-regularized logistic regression, results in better performance than all other models considered. These results suggest that GenePT’s literature-based embedding captures information relevant to gene and protein interactions; a promising future direction is to combine GenePT embeddings with protein embeddings learned from 3D structures or protein language models.

Finally, we delved into cell-type specific activations among the GenePT-derived gene programs within human immune tissue datasets through a “zero-shot” approach. We first constructed a similarity graph based on cosine similarities between the GenePT embeddings by placing an edge between two genes if the cosine similarity is larger than 0.9 and applied Leiden clustering to the resulting graph at a resolution of 20. Randomly sampled 20 gene programs comprising 10 or more genes are depicted in Figure 2(g). Here, we display the average expression levels of these gene programs, stratified by cell types. The observed selective activation of these programs aligns with established biological knowledge where the identified gene sets are known to be functionally distinct and are differentially expressed across different cell types (e.g., Gene set 8 comprising of IFI families and gene set 24 comprising of CDC families). These findings underscore that GenePT-inferred gene programs effectively capture biologically pertinent functional groups; additional results with different similarity thresholds can be found in Appendix B.4.

### 4.2 GenePT embeddings enable accurate predictions in chromatin dynamics and dosage sensitivity

In this section, we delve into specific biological tasks that predict the roles of genes in network dynamics with datasets curated from the literature by Theodoris et al. [1]: dosage-sensitive versus dosage-insensitive TFs, bivalent versus non-methylated genes, Lys4-only-methylated versus non-methylated genes, and long-versus short-range TFs. These tasks were used to demonstrate the utility of Geneformer. We assess the performance of GenePT and Gene2vec embeddings by five-fold cross-validated ROC-AUC with either an *ℓ*_2_ penalized logistic regression (LR) or a Random Forest (RF) classifier using default parameters from scikit-learn [45]. By contrast, Geneformer results, as reported in Theodoris et al. [1], are based on a fine-tuned transformer model. We also reported some variants of the GenePT framework: BioLinkBert embedding of the gene summaries; or GPT-3.5 embedding of only the gene names (without context or descriptions); and random embeddings matching the GenePT dimension (*d* = 1, 536). Table 1 illustrates that GenePT embeddings consistently achieve competitive results, sometimes even surpassing Geneformer, although the latter benefits from a substantial pre-training dataset and a more intricate classification head. Interestingly, GPT-3.5 embeddings of only gene names also show high accuracies in some tasks. This might be due to two aspects: 1. gene nomenclature attempts to designate functionally-related or homologous genes with similar symbols to enable grouping [46]; and 2. the underlying language model and tokenizer for GPT-3.5 might grasp the biological significance of these gene symbols due to extensive pretraining on scientific text [47]. Open-source embeddings like BioLinkBert and Gene2vec have slightly less competitive performance. As expected, random embeddings exhibit results similar to random guessing. The stark contrast in predictive performance between GenePT and random embeddings indicates that it’s unlikely that the GenePT performance is simply due to a large embedding dimension (*d* = 1, 536). In addition, since we used low-complexity, off-the-shelf *ℓ*_2_-regularized logistic regression and random forests, and reported results based on five-fold cross-validation, it is unlikely that the performance is due to model overfitting. In summary, these results underscore the potential of our versatile GenePT approach, which compares favourably with state-of-the-art deep learning models specifically crafted for single-cell RNA sequencing data.

**Table 1:**
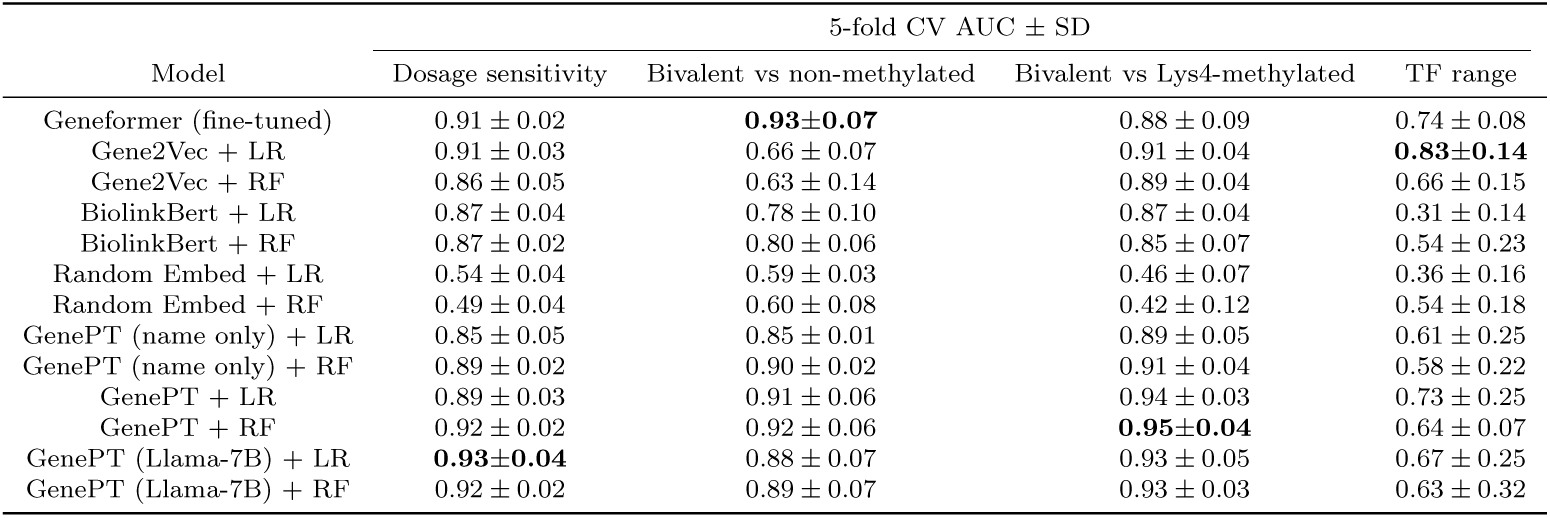
Cross-validated AUC for GenePT predictions versus alternative embeddings for downstream tasks of distinguishing (i) dosage-sensitive vs. insensitive transcription factors; (ii) bivalent versus non-methylated gene; (iii) bivalent versus Lys4-only methylated genes; and (iv) long-range versus short-range transcription factors (TFs). The performance for Geneformer is reproduced from Theodoris et al. [1] and is based on a fine-tuned sequence classification model. Here, random embed denotes an embedding identical in size to GenePT with entries drawn from i.i.d. *N* (0, 1). This serves as a “negative control” to ensure that signals in GenePT are not merely due to a larger embedding dimension. We use RF and LR to denote random forest and logistic regression models with default parameters in scikit-learn, respectively.

| Model | 5-fold CV AUC ± SD | | | |
| --- | --- | --- | --- | --- |
|  | Dosage sensitivity | Bivalent vs non-methylated | Bivalent vs Lys4-methylated | TF range |
| Geneformer (fine-tuned) | 0.91 ± 0.02 | **0.93±0.07** | 0.88 ± 0.09 | 0.74 ± 0.08 |
| Gene2Vec + LR | 0.91 ± 0.03 | 0.66 ± 0.07 | 0.91 ± 0.04 | **0.83±0.14** |
| Gene2Vec + RF | 0.86 ± 0.05 | 0.63 ± 0.14 | 0.89 ± 0.04 | 0.66 ± 0.15 |
| BiolinkBert + LR | 0.87 ± 0.04 | 0.78 ± 0.10 | 0.87 ± 0.04 | 0.31 ± 0.14 |
| BiolinkBert + RF | 0.87 ± 0.02 | 0.80 ± 0.06 | 0.85 ± 0.07 | 0.54 ± 0.23 |
| Random Embed + LR | 0.54 ± 0.04 | 0.59 ± 0.03 | 0.46 ± 0.07 | 0.36 ± 0.16 |
| Random Embed + RF | 0.49 ± 0.04 | 0.60 ± 0.08 | 0.42 ± 0.12 | 0.54 ± 0.18 |
| GenePT (name only) + LR | 0.85 ± 0.05 | 0.85 ± 0.01 | 0.89 ± 0.05 | 0.61 ± 0.25 |
| GenePT (name only) + RF | 0.89 ± 0.02 | 0.90 ± 0.02 | 0.91 ± 0.04 | 0.58 ± 0.22 |
| GenePT + LR | 0.89 ± 0.03 | 0.91 ± 0.06 | 0.94 ± 0.03 | 0.73 ± 0.25 |
| GenePT + RF | 0.92 ± 0.02 | 0.92 ± 0.06 | **0.95±0.04** | 0.64 ± 0.07 |
| GenePT (Llama-7B) + LR | **0.93±0.04** | 0.88 ± 0.07 | 0.93 ± 0.05 | 0.67 ± 0.25 |
| GenePT (Llama-7B) + RF | 0.92 ± 0.02 | 0.89 ± 0.07 | 0.93 ± 0.03 | 0.63 ± 0.32 |

Finally, it’s crucial to confirm that the promising results in Sections 4.1 and 4.2 are not simply the result of information leakage, such as test set data being included in the original NCBI gene summaries used as input for GenePT. We address these concerns in detail in Appendix B.3.

### 4.3 GenePT learns representations that reflect cell biology

In this section, we focus on determining the capacity of our cell embedding approaches, as depicted in Figure 1(b)–(c), in capturing the biology underpinning selected single-cell datasets. We sought to evaluate whether the GenePT embeddings are congruent with metadata annotations across six datasets representing cells from circulatory systems (Aorta and Artery), bone tissues (Bones, Myeloid), the Pancreas, and immune cells collected from healthy individuals and patients with Multiple Sclerosis.

We quantified the concordance between biological annotations (i.e., cell types, cancer types, donor ages) and *k*-means clustering labels inferred from: (i) pretrained Geneformer embeddings; (ii) pretrained scGPT embeddings; (iii) GenePT-w embeddings (as in Figure 1(b)); and (iv) GenePT-s embeddings (as in Figure 1(c)). We quantified the concordance using both AMI and ARI in Table 2. We see that latent representations via GenePT-s broadly outperformed both the GenePT-w and Geneformer embeddings in terms of AMI and ARI metrics and stayed competitive with the scGPT embeddings: across nine tasks, scGPT and GenePT each provides the most biological signal on five and four tasks, respectively. This demonstrates that GenePT cell embeddings capture biological variations comparable to two leading single-cell foundation models. An important caveat is that concordance with cell types and annotations is a limited measure of the utility of embedding, though it is widely used. We also included additional classification results for a cell type annotation task via a nearest neighbour approach on these datasets in Appendix C, which yields very similar findings that scGPT and GenePT-w are two of the best-performing methods in this setting, and both consistently outperform Geneformer in terms of prediction accuracy. Interestingly, a simple ensembling of the nearest neighbours retrieved by different embeddings (GenePT-w, GenePT-s, and scGPT) enhanced the predictive performance (see Table C4 in Appendix C). This suggests that natural language embeddings, such as GenePT, could provide complementary insights to existing expression-derived foundation models like scGPT in single-cell biology tasks.

**Table 2:** Assessing the Association Between Different Latent Cell Representations and Biological Annotations. This analysis involves datasets representing cells from circulatory systems (Aorta and Artery), bone tissues (Bones, Myeloid), the Pancreas, and immune cells collected from healthy individuals and patients with Multiple Sclerosis. We utilized pretrained Geneformer and scGPT embeddings for this task. The Adjusted Rand Index (ARI) and Adjusted Mutual Information (AMI) were computed to compare the labels derived from *k*-means clustering with the true annotations of the original samples (higher values indicate better alignment); the Average Silhouette Width (ASW) was calculated using the true annotations of original samples to assess the cohesion and separation of the clusters.

| Dataset | Annotation | Geneformer |  |  | scGPT |  |  | GenePT-w |  |  | GenePT-s |  |  |
| --- | --- | --- | --- | --- | --- | --- | --- | --- | --- | --- | --- | --- | --- |
|  |  | ARI | AMI | ASW | ARI | AMI | ASW | ARI | AMI | ASW | ARI | AMI | ASW |
| Aorta | Phenotype | 0.10 | 0.12 | -0.005 | **0.12** | **0.12** | 0.01 | 0.09 | **0.12** | -0.04 | **0.12** | 0.11 | **0.02** |
|  | Cell type | 0.21 | 0.31 | -0.04 | 0.47 | **0.64** | 0.18 | **0.54** | 0.60 | 0.03 | 0.31 | 0.47 | 0.04 |
| Artery | Cell type | 0.39 | 0.59 | 0.10 | 0.36 | 0.59 | 0.15 | **0.42** | **0.67** | **0.16** | 0.36 | 0.56 | 0.06 |
| Bones | Cell type | 0.09 | 0.16 | -0.01 | 0.12 | 0.21 | -0.01 | **0.21** | **0.29** | **0.02** | 0.17 | 0.28 | 0.003 |
| Myeloid | Cancer type | 0.16 | 0.18 | 0.03 | **0.27** | **0.29** | **0.08** | 0.25 | 0.27 | 0.02 | 0.17 | 0.17 | 0.06 |
|  | Cell type | 0.19 | 0.29 | -0.02 | **0.44** | **0.53** | **0.13** | 0.21 | 0.28 | 0.001 | 0.30 | 0.41 | 0.03 |
| Pancreas | Cell type | 0.04 | 0.11 | -0.09 | 0.21 | 0.41 | 0.05 | **0.49** | **0.69** | **0.15** | 0.30 | 0.50 | 0.10 |
| Multiple Sclerosis | Age | 0.04 | 0.11 | -0.1 | 0.04 | 0.11 | -0.06 | **0.07** | **0.13** | -0.07 | 0.06 | 0.12 | **-0.03** |
|  | Cell type | 0.21 | 0.35 | -0.05 | **0.25** | **0.44** | **0.04** | 0.17 | 0.32 | -0.02 | 0.19 | 0.35 | 0.002 |

### 4.4 GenePT embedding removes batch effect while preserving underlying biology

We next assess whether GenePT embeddings are robust to batch-dependent technical artifacts such as patient variability. We compared the performance of GenePT with pretrained Geneformer and scGPT using a 10% random sample from a cardiomyocyte dataset by Chaffin et al. [44] and a 20% random sample from the Aorta dataset consisting of cells in healthy and dilated aortas [39], both of which were used to demonstrate the utility of Geneformer.

In the cardiomyocyte dataset, the scientific question was to distinguish cardiomyocytes in non-failing hearts from those in hypertrophic or dilated cardiomyopathy samples. Notably, the original data exhibited significant patient batch effects (see Figure D8(b) in the Appendix). We performed the following analysis to quantify the patient-level batch effects: (i) we first project the data (either the original RNA-seq or one of the pretrained embeddings) into the top 50 principal components; (ii) we then applied *k*-means clustering with *k* = 42, which is the number of distinct patients; (iii) we compute adjusted Rand index (ARI) between the cell clusters and patient clusters. Higher ARI values indicate more patient-level batch effects. The original scRNA-seq data has a high ARI of 0.33, suggesting strong batch effects. Using the GenePT-s, Geneformer, and scGPT, the ARI dropped to 0.07, 0.01, and 0.01 respectively, showing that these embeddings are robust to batch effects.

In addition to reducing batch effects, we also investigated whether these embeddings could preserve the underlying disease phenotype (i.e., non-failing versus cardiomyopathy) of the patients from whom the cells were collected. To this end, we randomly split the cardiomyocytes into 80%/20% train/test sets and evaluated the predictive performance using the *ℓ*_2_-regularized logistic regression on top of the following pre-trained embeddings: (i) GenePT-s, (ii) scGPT, and (iii) Geneformer. Overall, GenePT-s and scGPT achieve nearly identical performance on the held-out test set (88% accuracy, 88% precision, and 88% recall for both embeddings for predicting disease label), whereas the performance for pretrained Geneformer trailed behind (71% accuracy, 72% precision, and 71% recall).

Next, we conducted a similar study with the Aorta dataset, collected over 11 patients (eight patients with Ascending thoracic aortic aneurysm (ATAA) and three control subjects; the eight ATAA patients are further divided into three different phenotypes: ascending only, ascending with descending thoracic aortic aneurysm, and ascending with root aneurysm). We demonstrate the use of GenePT on a random 20% sample of the original Aorta dataset. In Figure 3, we display the original data (top panel) and GenePT-s embeddings (bottom panel) using UMAP, colored by patient phenotype (left panel), annotated cell types (middle panel), and patient identity (right panel). While the original data was highly influenced by patient batch effect (see Figure 3(c)) and displayed distinct clusters for identical cell types (e.g., T cells and Mono/Maph/Dend cells in Figure 3(b)), GenePT-s embeddings clustered primarily by cell types (Figure 3(e)) as well as disease phenotype (Figure 3(d)). In particular, GenePT-s embeddings were able to distinguish the phenotype of ascending only aortic aneurysm (green points in Figure 3(d)), a different phenotype than aortic aneurysm that includes the root (purple points in Figure 3(d)).

**Fig. 3:**
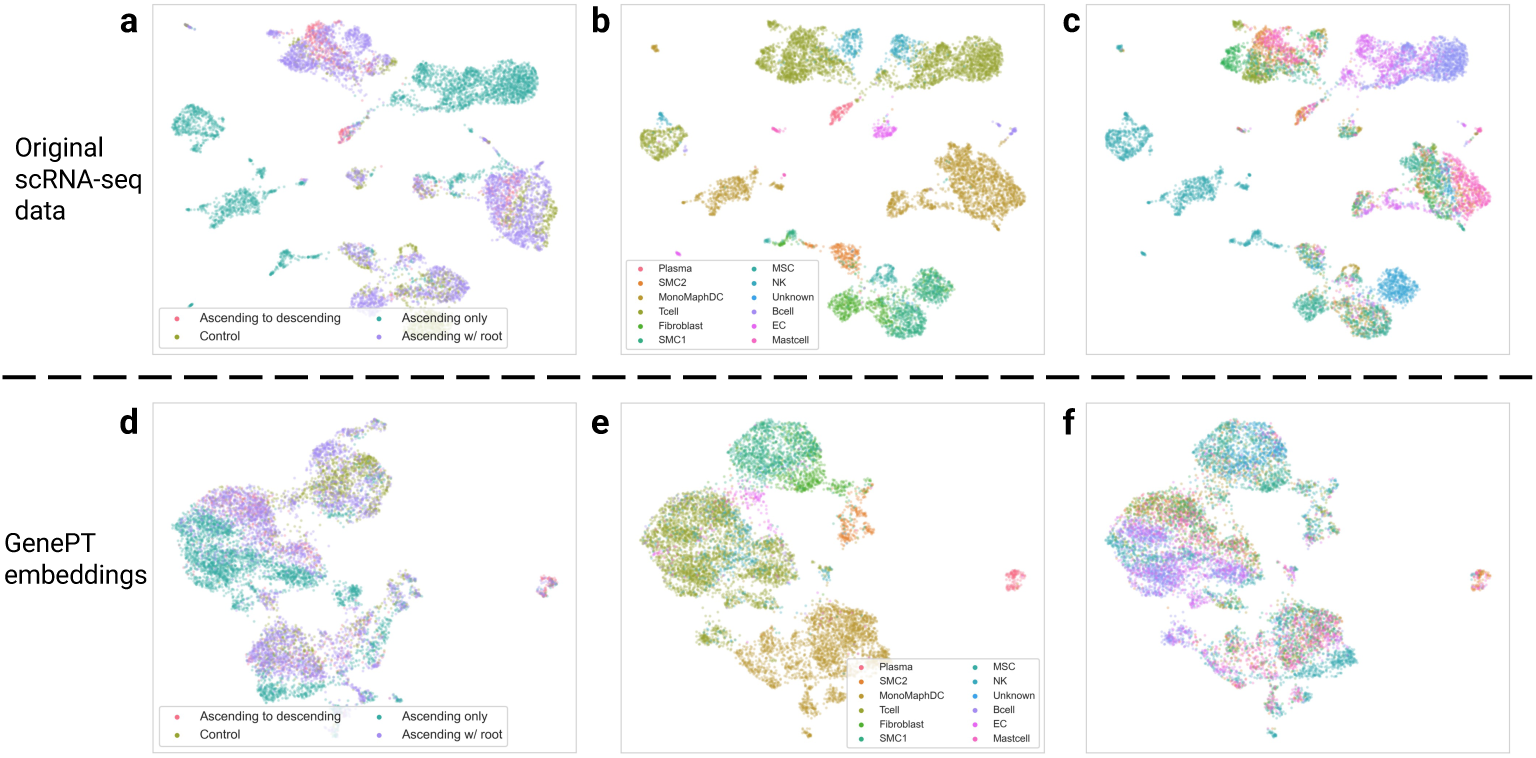
**(a)** UMAP visualization of the subsampled Aorta dataset, coloured by disease phenotype (three different disease phenotypes: ascending only, ascending with descending thoracic aortic aneurysm, and ascending with root aneurysm; one control phenotype comprising patients with healthy hearts after transplant) provided in the original study [39]. **(b)** Same as **(a)**, but colored by cell types annotated by the original study [39]. **(c)** Same as **(a)**, but colored by patient id. **(d)** UMAP visualization of GenePT-s embeddings of the same set of cells as **(a)**, coloured by disease phenotype. **(e)** Same as **(d)**, but colored by cell types. **(f)** Same as **(d)**, but colored by patient identity. further corroborated by training a logistic regression model to predict the phenotype: on the randomly held-out 20% test set, GenePT-s yields an accuracy of 73% (68% precision, 74% recall), similar to that of scGPT (75% accuracy, 75% precision, 75% recall) and moderately better than Geneformer (69% accuracy, 68% precision, 69% recall).

We repeated the clustering analysis above on the Aorta dataset to get a more quantitative measure of patient-level batch effects. The ARI between patient labels and the estimated *k*-means clusters (*k* = 11) on the original scRNA-seq data is 0.24 versus 0.11, 0.10, and 0.18 when using Geneformer, GenePT-s, and scGPT respectively. We also evaluated the agreement between the phenotype labels (three ATAA subtypes and one control) and the clusters derived from embeddings and original scRNA-seq data. The resulting ARIs are 0.12, 0.11, 0.12, and 0.12 for Geneformer embeddings, GenePT-s embeddings, scGPT embeddings, and scRNA-seq data, respectively. These findings suggest that GenePT-s, Geneformer, and scGPT all exhibit some degree of robustness against batch effects while preserving information on the disease phenotype. This is

## 5 Discussion

With the advance of technologies to measure genetic and cellular functionalities, enhancing our understanding of the underlying biology through latent embedding representations has attracted much interest. In this work, we introduced GenePT, a simple yet effective approach that leverages GPT-3.5 to represent genes and cells by utilizing their text summaries and ranked expression values, respectively. Across various contexts, including discerning gene functionality groups and predicting gene-gene interactions, this straightforward approach proves to be very effective even compared to state-of-the-art foundational models trained on large-scale single-cell transcriptomics data. Our work underscores the potential of complementing those specially crafted foundational models with a simple, natural language-guided representation, which could be substantially more resource and data-efficient.

It is important to note the limitations in our work, primarily because the current GenePT framework only makes use of available gene summaries and descriptions. This may overlook the intricacies of lesser-known functionalities not documented in databases like NCBI. Furthermore, unlike the embeddings trained on expression data, GenePT embeddings might not be optimal for specific tissues and cell types. This might pose challenges in capturing the dynamic and context-dependent roles of genes and cells within those settings. Lastly, the effectiveness of the embeddings is inherently constrained by the language models employed, i.e., GPT-3.5. Fine-tuning the language models could further enhance understanding of the domain-specific language prevalent in genomics.

Several promising pathways lie ahead for future research. First, extending the current GenePT approach to be more dynamic and context-dependent — such as via fine-tuning models and incorporating tissue, disease, and marker-gene-specific information — could enhance its utility in real-world applications. Additionally, investigating ways to integrate different embeddings across different single-cell foundation models, as well as leveraging recent developments in dimension reduction techniques [48] to obtain more compact representations, would be an important realm of future work. Moreover, it’s natural to investigate the performance of GenePT in additional downstream tasks, such as perturbation predictions and drug-gene interactions. Lastly, while this paper primarily focuses on gene and cell embeddings, it would be of great interest to explore whether the approach of leveraging the natural language descriptions with LLMs embedding could be applied to other biological domains and challenges, such as protein sequence modeling [49] and Genome-Wide Association Studies [50].

## Supplementary information

Supplementary information contains additional details on the methods and results, as well as Table B2, Figure D8, and Figure 3.

## Data availability

All datasets used in the study have been previously published with pointers provided at https://github.com/yiqunchen/GenePT.

## Code availability

GenePT is available at https://github.com/yiqunchen/GenePT.

## Acknowledgments

YC is supported by a Stanford Data Science Postdoctoral Fellowship. JZ is supported by the National Science Foundation (CCF 1763191 and CAREER 1942926), the US National Institutes of Health (P30AG059307 and U01MH098953) and grants from the Silicon Valley Foundation and the Chan-Zuckerberg Initiative.

## Appendix A Assessing GenePT sensitivity to variations in gene summary inputs

In this section, we present additional details on the example gene summary inputs to GenePT, as well as sensitivity analyses using three different levels of content input for gene summaries: gene names only, gene names and gene summaries only, and all summary card information.

As a concrete example, for the gene CD24, the three levels correspond to:

1. CD24
2. Gene Name CD24 Summary This gene encodes a sialoglycoprotein that is expressed on mature granulocytes and B cells and modulates growth and differentiation signals to these cells. The precursor protein is cleaved to a short 32 amino acid mature peptide which is anchored via a glycosyl phosphatidylinositol (GPI) link to the cell surface. This gene was missing from previous genome assemblies, but is properly located on chromosome 6. Non-transcribed pseudogenes have been designated on chromosomes 1, 15, 20, and Y. Alternative splicing results in multiple transcript variants. Expression Biased expression in thyroid (RPKM 586.8), esophagus (RPKM 431.3) and 12 other tissues
3. CD24 Official Full Name CD24 molecule Primary source HGNC:HGNC:1645 See related Ensembl:ENSG00000272398 MIM:600074; AllianceGenome:HGNC:1645 Gene type protein coding RefSeq status REVIEWED Also known as CD24A Summary This gene encodes a sialoglycoprotein that is expressed on mature granulocytes and B cells and modulates growth and differentiation signals to these cells. The precursor protein is cleaved to a short 32 amino acid mature peptide which is anchored via a glycosyl phosphatidylinositol (GPI) link to the cell surface. This gene was missing from previous genome assemblies, but is properly located on chromosome 6. Non-transcribed pseudogenes have been designated on chromosomes 1, 15, 20, and Y. Alternative splicing results in multiple transcript variants. Expression Biased expression in thyroid (RPKM 586.8), esophagus (RPKM 431.3) and 12 other tissues See more Orthologs mouse all.

In the most comprehensive input (item 3, as previously detailed), we observe that the information provided for a specific gene typically includes two distinct types of content: (i) various names, symbols, RefSeq status, and orthologs, as well as (ii) detailed summaries of gene types, functions, and notable expression levels, which are concisely summarized in item 2 of the list.

To assess the sensitivity of input text to GenePT, we have provided results for genelevel classification and gene-gene interaction inference below (refer to Table A1 and Figure A1). Our results indicate that most of the biological information, as expected, is encoded in the gene summary from the NCBI gene dataset. In addition, adding gene function summaries substantially improves the performance in the biological tasks considered, compared to using gene names alone. We expect that our pipeline is robust to different levels of input cleaning as long as the gene function summaries are included.

**Table A1:** Cross-validated AUC for GenePT predictions versus alternative embeddings for downstream tasks of distinguishing (i) dosage-sensitive vs. insensitive transcription factors; (ii) bivalent versus non-methylated gene; (iii) bivalent versus Lys4-only methylated genes; and (iv) long-range versus short-range transcription factors (TFs). We use RF and LR to denote random forest and logistic regression models with default parameters in the scikit-learn package [45], respectively.

| Model | 5-fold CV AUC ± SD | | | |
| --- | --- | --- | --- | --- |
|  | Dosage sensitivity | Bivalent vs non-methylated | Bivalent vs Lys4-methylated | TF range |
| GenePT (name only) + LR | 0.85 ± 0.05 | 0.85 ± 0.01 | 0.89 ± 0.05 | 0.61 ± 0.25 |
| GenePT (name only) + RF | 0.89 ± 0.02 | 0.90 ± 0.02 | 0.95 ± 0.04 | 0.64 ± 0.07 |
| GenePT (summary only) + LR | 0.87 ± 0.04 | 0.91 ± 0.05 | 0.95 ± 0.03 | 0.74 ± 0.14 |
| GenePT (summary only) + RF | 0.89 ± 0.04 | 0.91 ± 0.07 | 0.95 ± 0.04 | 0.86 ± 0.14 |
| GenePT + LR | 0.89 ± 0.03 | 0.91 ± 0.06 | 0.94 ± 0.03 | 0.73 ± 0.25 |
| GenePT + RF | 0.92 ± 0.02 | 0.92 ± 0.06 | 0.95 ± 0.04 | 0.64 ± 0.07 |

**Fig. A1:**
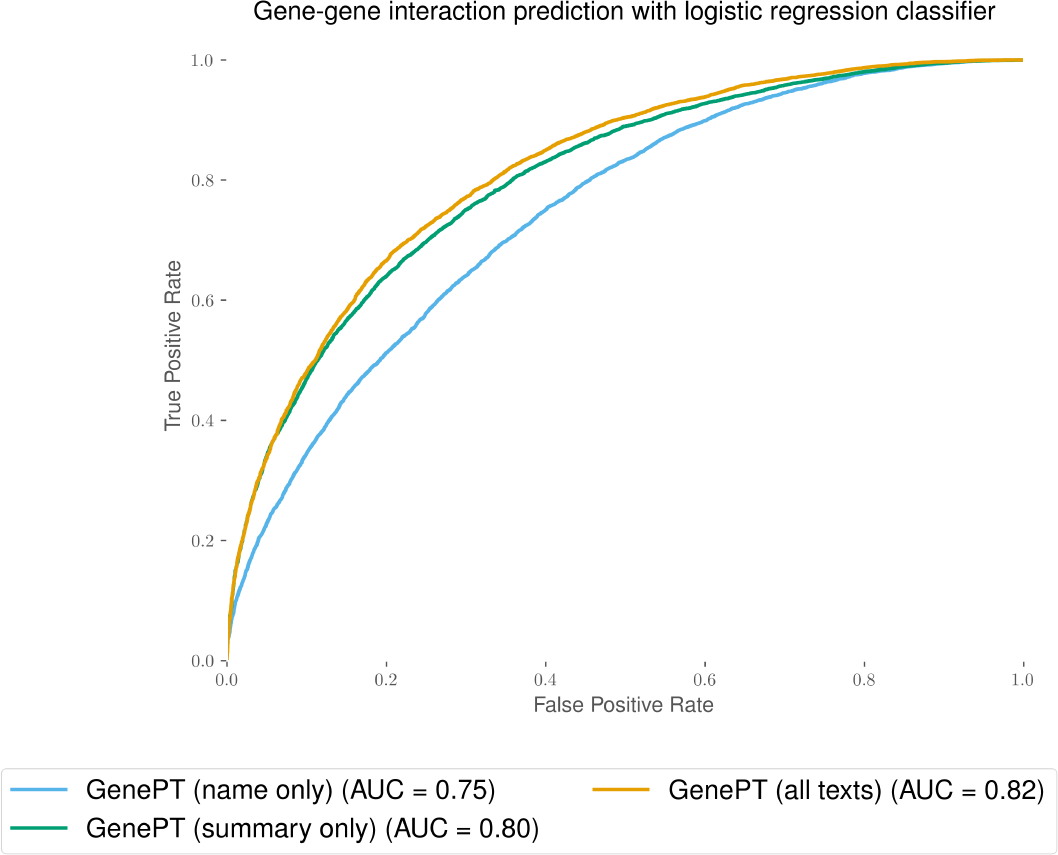
Test set prediction performance using text-embedding-ada-002 embedding model on different text input (name only, cleaned gene summary only, all gene information) from NCBI dataset. The gene-gene interaction benchmark dataset used here was derived from GEO expression data [18].

## Appendix B Additional results for the gene level functionality and property predictions

### >B.1 Gene functionality

In Figure 2(b), we display the results from a prediction task that leverages GenePT embedding to predict the 15 most prevalent gene functional class, as detailed in Table B2 below. For baseline comparison, we also applied LR classification using Gene2vec embeddings on the subset of the genes that had corresponding Gene2vec embeddings (21,000 genes). This yielded a five-fold cross-validated accuracy of 0.86 (SD: 0.03). On the same subset of 21, 000 genes, GenePT achieved a considerably higher average accuracy of 0.95 (SD:0.05).

**Table B2:** Gene functional classes used in the prediction task of Figure 2(b).

| Type | Count | Percentage (%) |
| --- | --- | --- |
| Protein coding | 20,184 | 71.6 |
| pseudogene | 3,725 | 13.2 |
| miRNA | 2,061 | 7.3 |
| snRNA | 1,793 | 6.4 |
| misc RNA | 1,043 | 3.7 |
| lncRNA | 724 | 2.6 |
| snoRNA | 692 | 2.5 |
| antisense | 592 | 2.1 |
| rRNA | 512 | 1.8 |
| lincRNA | 499 | 1.8 |
| processed transcript | 248 | 0.9 |
| IG V gene | 97 | 0.3 |
| sense intronic | 84 | 0.3 |
| IG V pseudogene | 79 | 0.3 |
| TR J gene | 76 | 0.3 |

### B.2 Protein-protein interaction

In Figure B2, we display the precision-recall curve for the three protein-protein interaction prediction tasks (see Section 3 for datasets descriptions and Figure 2(d)–(f) for ROC curves).

### B.3 Addressing potential information leakage

It’s important to ensure that the promising results in Section 4 are not merely due to the test set being represented in the original NCBI gene summaries used as input for GenePT gene embeddings. We address the concerns of information leakage by using a temporal split for gene functionality prediction and quantifying the extent of information leakage for gene-gene and protein-protein interaction predictions.

1. Temporal Split: In terms of the temporal split, we discovered that the gene functionality data was released by Theodoris et al. [1], in June 2023 at https://huggingface.co/datasets/ctheodoris/Genecorpus-30M/tree/main/example input files. Conversely, the NCBI gene information does not contain information on these specific data, and the embedding model was released in December 2022. Therefore, the information leakage for the gene property prediction should be minimal. Regarding gene functionality, we intended this as a sanity check to ensure that our embeddings contain first-order information, and we have adjusted our original statement accordingly.
2. Content Overlap: We have also quantified the extent to which explicit gene-gene interactions and protein-protein interactions are mentioned in the NCBI gene summaries (see Table B3). This ensures that we are not merely memorizing training data. In this context, we quantified, for gene-gene interaction and protein-protein interaction datasets, the number of explicitly mentioned interaction pairs. We note that the number of present pairs is quite small across these datasets. In particular, except for the Lit-BM PPI dataset, the small percentage (less than 1%) of leakage in the other three datasets had a minimal impact on the classification result.

**Fig. B2:**
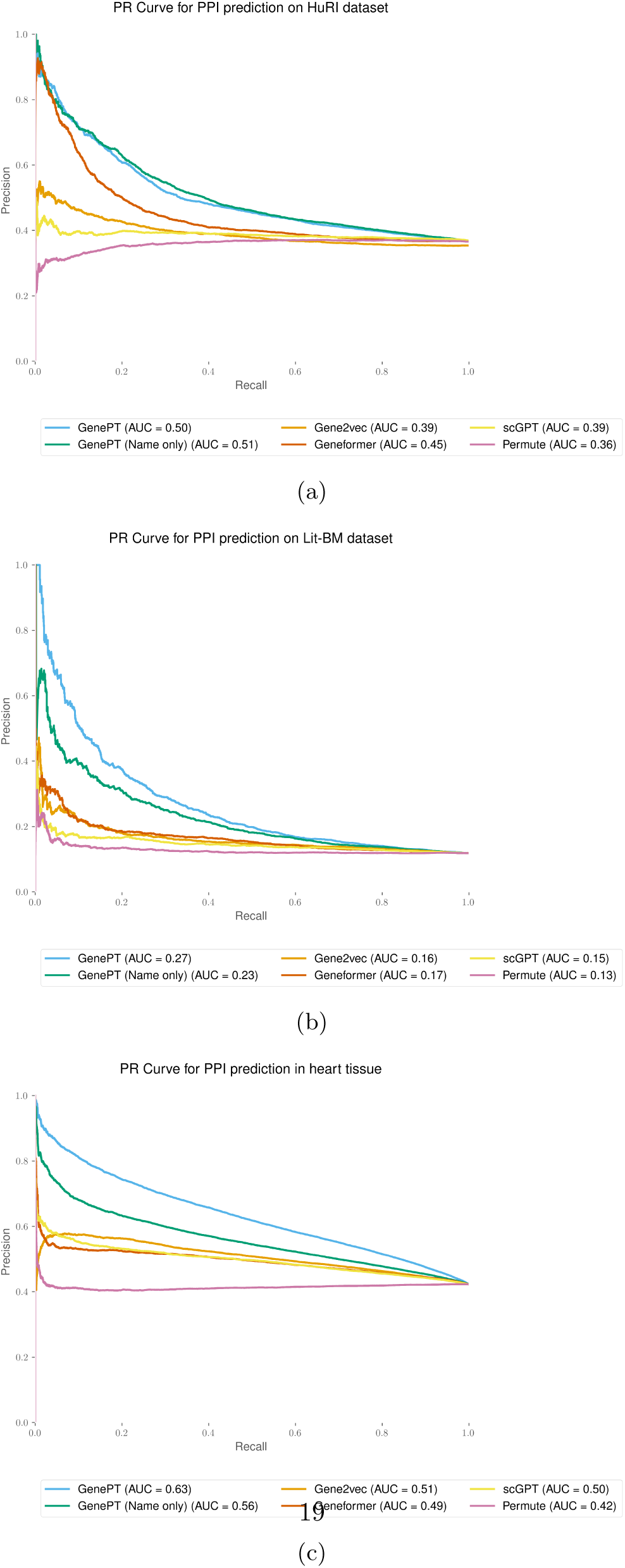
GenePT gene embeddings lead to the best test set prediction performance for all protein-protein interaction datasets considered, measured by the PrecisionRecall curve for high-quality binary literature datasets [34] (panel (a)), human binary protein interactions dataset [33] (panel (b)), and human heart tissue protein-protein functional interactions [35].

Additionally, to quantify the impact of data leakage in the Lit-BM PPI dataset (approximately 4%), we removed the positive pairs present in the NCBI summary and evaluated the performance again. We observed that there is no meaningful difference between the performance with and without the pairs that were mentioned in the NCBI summary. Therefore, this establishes that our GenePT approach was able to encode more information than mere memorization.

**Table B3:** Quantifying potential information leakage in gene-gene interaction and protein-protein interaction datasets.

| Dataset | Positive pairs | Pairs present in NCBI summary |
| --- | --- | --- |
| GGI [18] | 141,130 | 310 |
| HuRI PPI [33] | 53,548 | 583 |
| Lit-BM PPI [34] | 12,234 | 524 |
| Heart tissue PPI [35] | 303,932 | 469 |

**Fig. B3:**
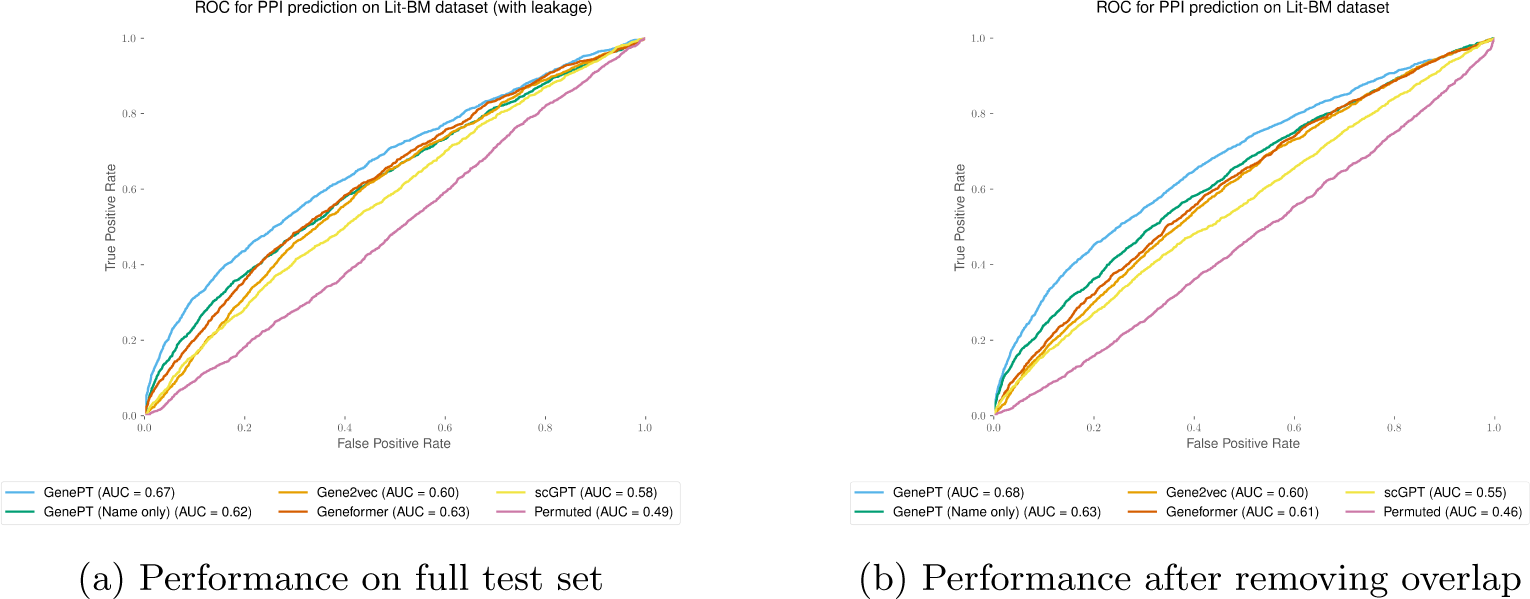
Test set prediction performance using GenePT gene embeddings on the Lit-BM protein-protein interaction datasets [34] before (panel (a)) and after (panel (b)) removing the mentioned pairs of interacting proteins in the NCBI gene summary database.

**Fig. B4:**
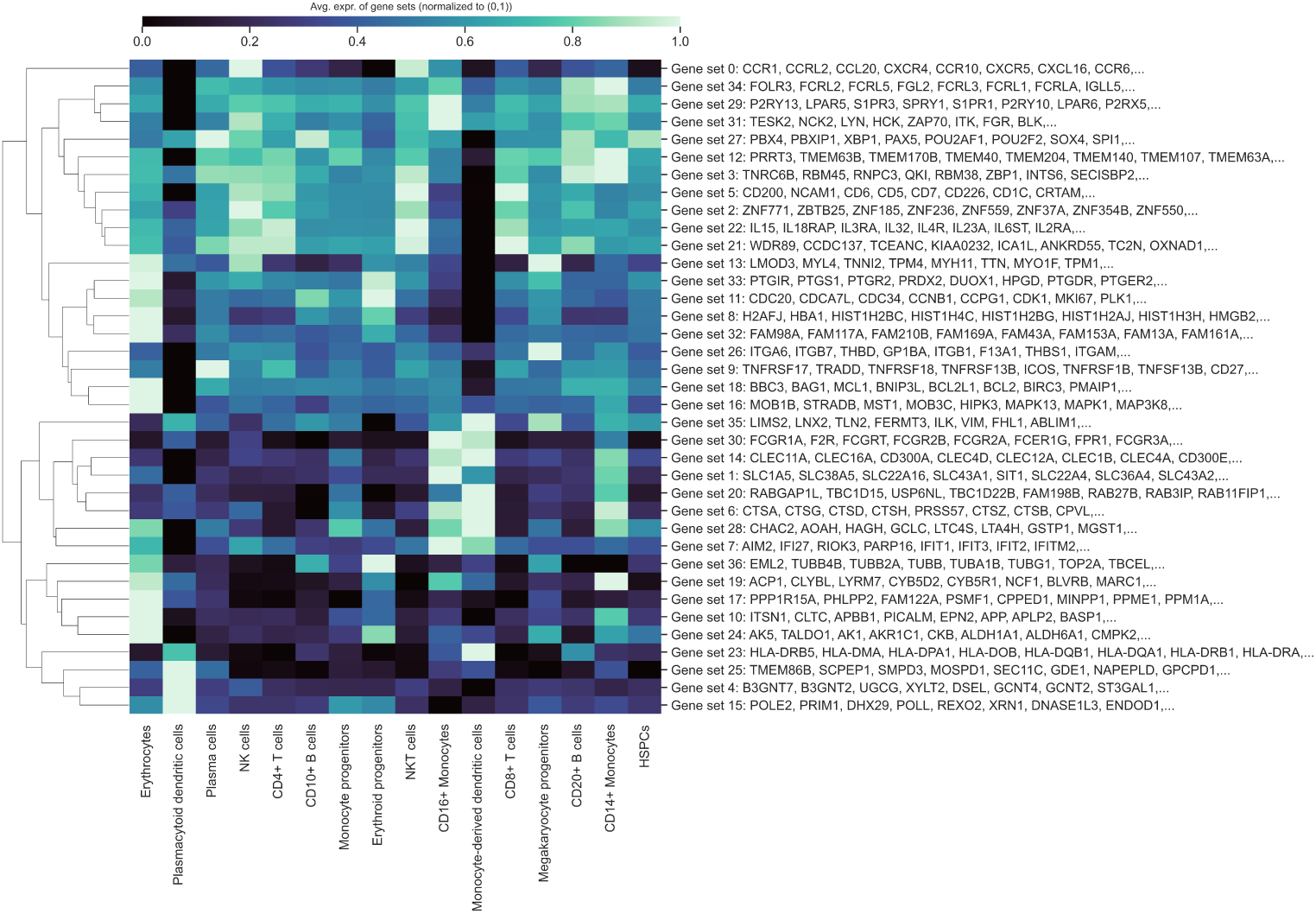
Cell-type specific activation among GenePT-embeddings-extracted gene programs (*>*10 in size) using similarity threshold 0.9 in a human immune tissue dataset [37].

### >B.4 Additional results on gene program identification

In this section, we provide additional results on the gene programs we identified in Figure 2(g). Specifically, we present gene programs comprising more than 10 genes at two different thresholds: 0.9 (the superset from which programs in Figure 2(g) were sampled) and 0.7, shown in Figures B4 and B5, respectively. We observe that the number and composition of different gene programs, as well as the overall cell-type-specific expression patterns of the identified programs, are similar across these two thresholds. This similarity indicates that while the optimal threshold may depend on the specific datasets’ downstream use, identifying programs using GenePT provides a stable method to group biologically relevant genes in a general context.

### >B.5 Experiments with tissue-dependent gene emebddings

In this section, we explore the potential utility of creating context-dependent embeddings by providing context-dependent gene descriptions. We experimented with this idea by prompting GPT-4 to generate tissue-specific contexts and then obtaining new GenePT gene embeddings based on these tissue-specific contexts.

Specifically, we give GPT-4 (model gpt-4-1106-preview) the following prompt:

**Fig. B5:**
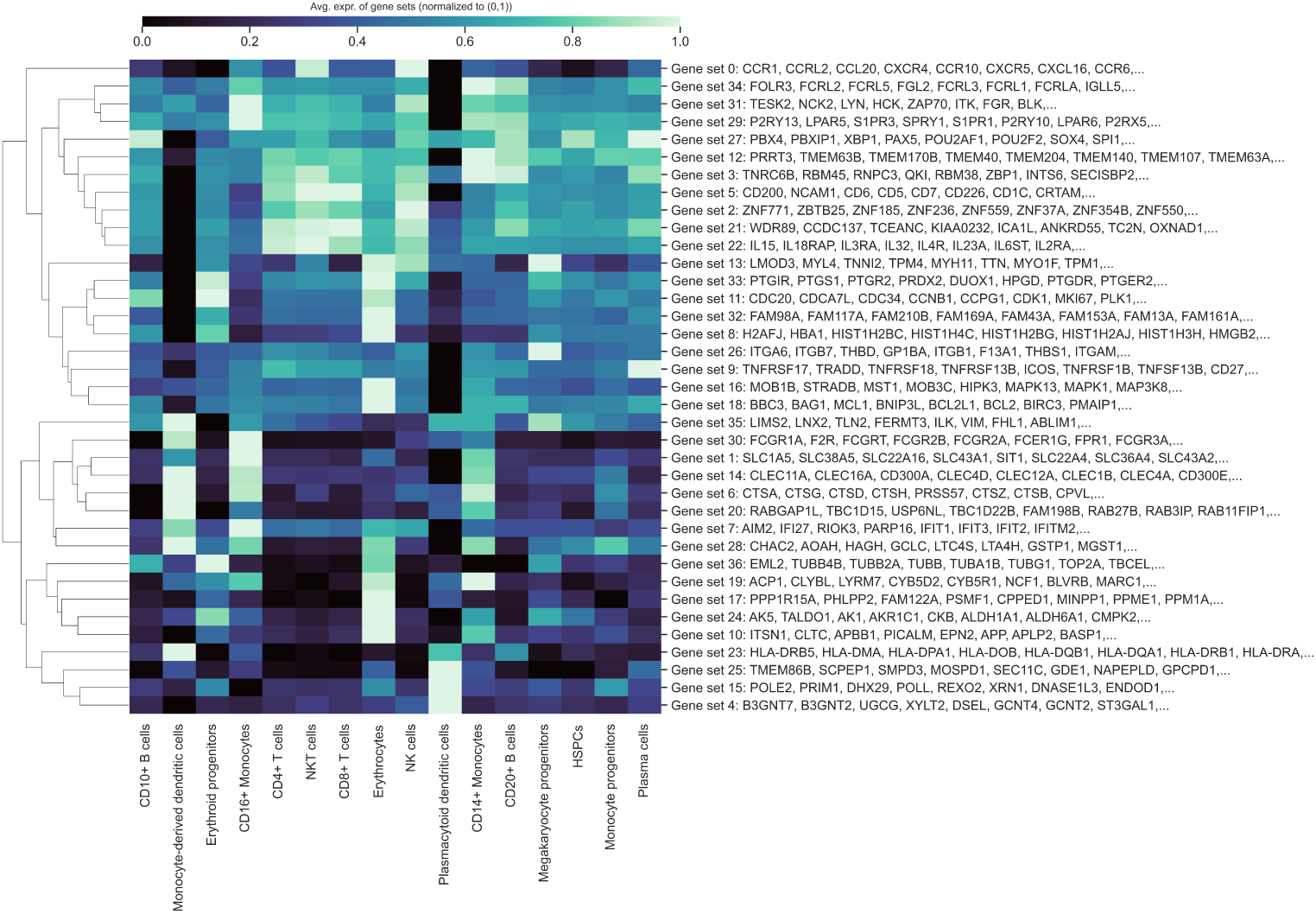
Cell-type specific activation among GenePT-embeddings-extracted gene programs (*>*10 in size) using similarity threshold 0.7 in a human immune tissue dataset [37].

Rewrite (the resulting summary should be approximately similar in length) the following summary of **gene X** to emphasize their functionality in the following different tissues or contexts: **tissue name**, **NCBI gene summary for gene X**.

We then concatenate the gene summary rewriting we get for a gene with the original NCBI summary and use the resulting text embedding as the tissue-specific gene embedding. We demonstrate in two protein-protein interaction studies that these context-specific embeddings were able to boost the prediction performance (see Figures B6 and B7.

## Appendix C Cell type annotation results

In this section, we also consider the cell type annotation task, where the primary aim is to predict annotated cell type labels based on the input cell representation. This annotation step is critical in single-cell analysis, as accurately distinguishing various cell populations within sequenced tissues can significantly enrich downstream biological insights. Mirroring the experimental design in the scGPT paper [2], we evaluated different embeddings’ efficacy for cell-type annotation using a 10-nearest-neighbor classifier on datasets representing cells from circulatory systems (Aorta and Artery), bone tissues (Bones, Myeloid), the Pancreas, and immune cells collected from healthy individuals and patients with Multiple Sclerosis. We report the test set classification accuracy by applying a 10-nearest neighbour classifier on various pretrained embeddings in Table C4 and note that GenePT embeddings held the ground against pretrained scGPT embeddings and outperformed the pretrained Geneformer embeddings. For the Aorta dataset, we used a random 80%/20% train/test split. Furthermore, we explored an ensemble approach that aggregates the 10 nearest neighbours from GenePT-w, GenePT-s, and scGPT, resulting in 30 predictions for each cell. This method demonstrated enhanced performance across various datasets and metrics. This indicates that literature-based natural language embeddings, such as GenePT-s, and expression-profile-derived embeddings like scGPT, provide complementary insights in single-cell biology tasks.

**Fig. B6:**
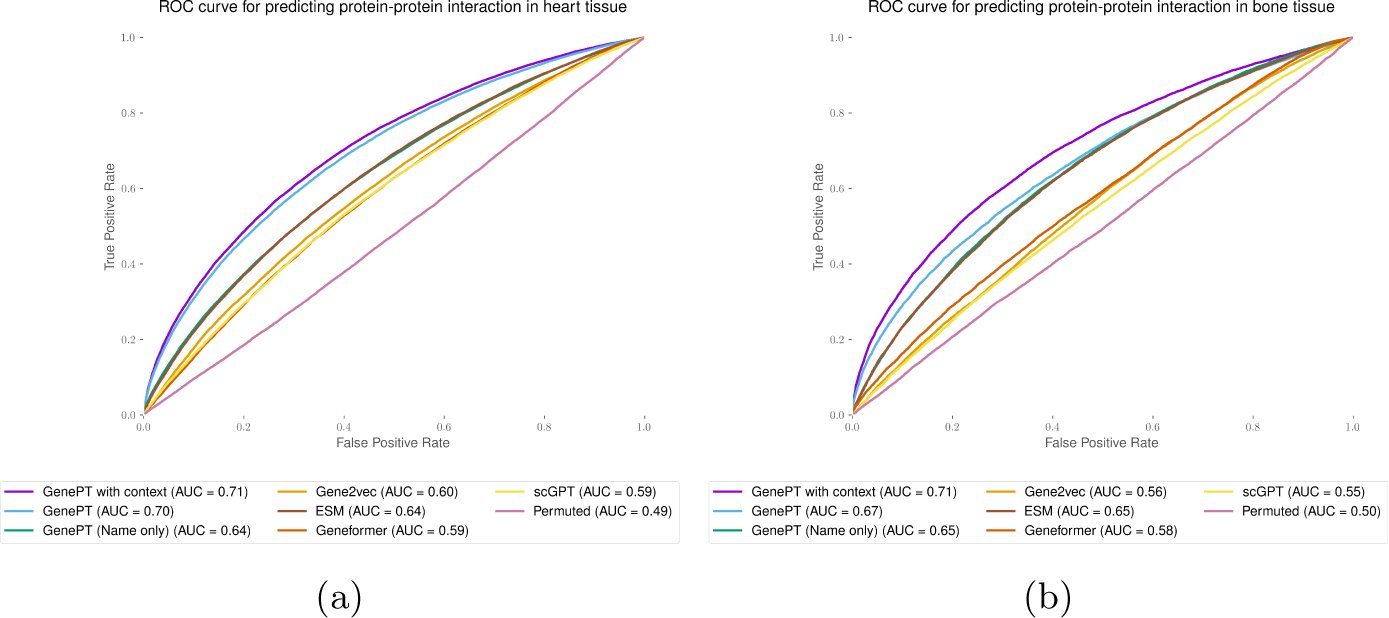
Context-enriched gene embeddings lead to better test set prediction performance, measured by the ROC curve for the heart (panel (a)) and bone (panel (b)) tissue protein-protein interaction dataset [35].

**Fig. B7:**
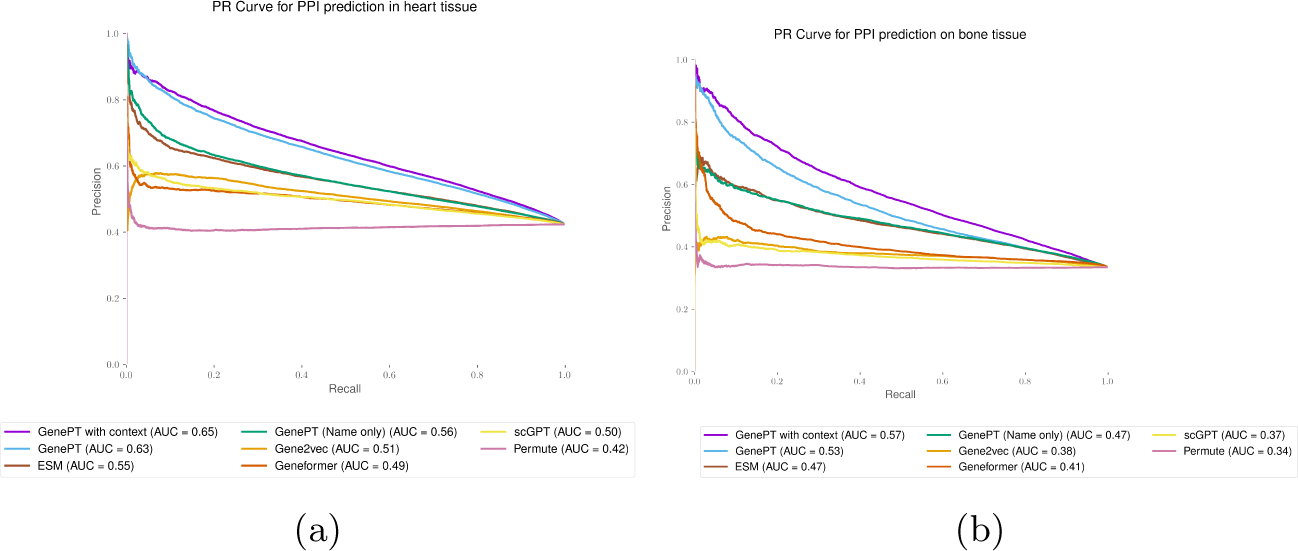
Context-enriched gene embeddings lead to better test set prediction performance, measured by the Precision-Recall curve for the heart (panel (a)) and bone (panel (b)) tissue protein-protein interaction dataset [35].

**Table C4:**
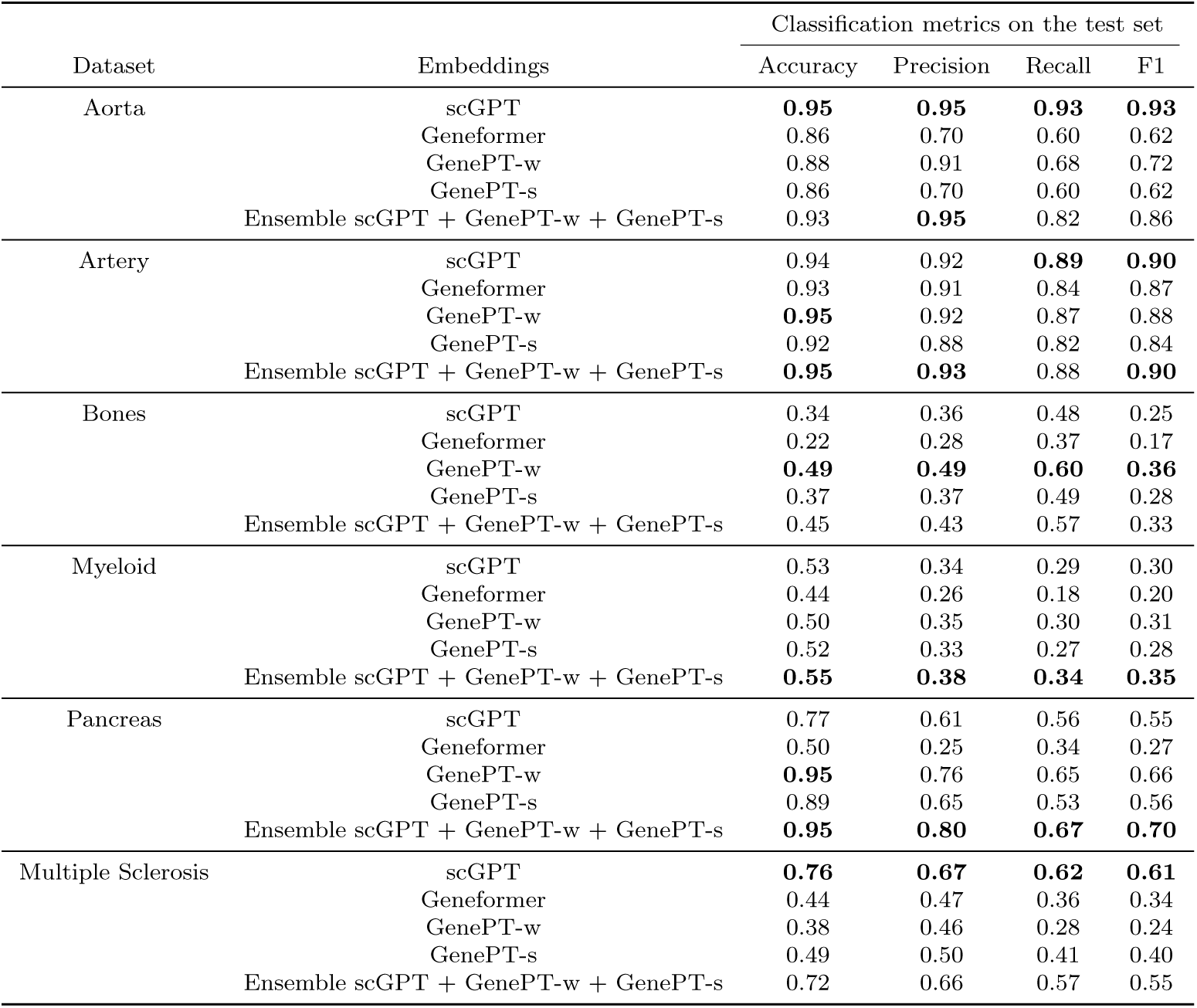
Test set performance on cell-type annotation tasks. This analysis involves datasets representing cells from circulatory systems (Aorta and Artery), bone tissues (Bones, Myeloid), the Pancreas, and immune cells collected from healthy individuals and patients with Multiple Sclerosis. Reported metrics include accuracy, precision, recall, and F1 (Macro-weighted), and are based on applying 10-nearest neighbour classifiers (using cosine similarity as the distance metric) on pretrained emebddings from scGPT, Geneformer, GenePT-w, and GenePT-s. We also report the performance of ensembling the nearest neighbours retrieved by scGPT, GenePT-w, and GenePT-s.

| Dataset | Embeddings | Classification metrics on the test set |  |  |  |
| --- | --- | --- | --- | --- | --- |
|  |  | Accuracy | Precision | Recall | F1 |
| Aorta | scGPT | **0.95** | **0.95** | **0.93** | **0.93** |
|  | Geneformer | 0.86 | 0.70 | 0.60 | 0.62 |
|  | GenePT-w | 0.88 | 0.91 | 0.68 | 0.72 |
|  | GenePT-s | 0.86 | 0.70 | 0.60 | 0.62 |
|  | Ensemble scGPT + GenePT-w + GenePT-s | 0.93 | **0.95** | 0.82 | 0.86 |
| Artery | scGPT | 0.94 | 0.92 | **0.89** | **0.90** |
|  | Geneformer | 0.93 | 0.91 | 0.84 | 0.87 |
|  | GenePT-w | **0.95** | 0.92 | 0.87 | 0.88 |
|  | GenePT-s | 0.92 | 0.88 | 0.82 | 0.84 |
|  | Ensemble scGPT + GenePT-w + GenePT-s | **0.95** | **0.93** | 0.88 | **0.90** |
| Bones | scGPT | 0.34 | 0.36 | 0.48 | 0.25 |
|  | Geneformer | 0.22 | 0.28 | 0.37 | 0.17 |
|  | GenePT-w | **0.49** | **0.49** | **0.60** | **0.36** |
|  | GenePT-s | 0.37 | 0.37 | 0.49 | 0.28 |
|  | Ensemble scGPT + GenePT-w + GenePT-s | 0.45 | 0.43 | 0.57 | 0.33 |
| Myeloid | scGPT | 0.53 | 0.34 | 0.29 | 0.30 |
|  | Geneformer | 0.44 | 0.26 | 0.18 | 0.20 |
|  | GenePT-w | 0.50 | 0.35 | 0.30 | 0.31 |
|  | GenePT-s | 0.52 | 0.33 | 0.27 | 0.28 |
|  | Ensemble scGPT + GenePT-w + GenePT-s | **0.55** | **0.38** | **0.34** | **0.35** |
| Pancreas | scGPT | 0.77 | 0.61 | 0.56 | 0.55 |
|  | Geneformer | 0.50 | 0.25 | 0.34 | 0.27 |
|  | GenePT-w | **0.95** | 0.76 | 0.65 | 0.66 |
|  | GenePT-s | 0.89 | 0.65 | 0.53 | 0.56 |
|  | Ensemble scGPT + GenePT-w + GenePT-s | **0.95** | **0.80** | **0.67** | **0.70** |
| Multiple Sclerosis | scGPT | **0.76** | **0.67** | **0.62** | **0.61** |
|  | Geneformer | 0.44 | 0.47 | 0.36 | 0.34 |
|  | GenePT-w | 0.38 | 0.46 | 0.28 | 0.24 |
|  | GenePT-s | 0.49 | 0.50 | 0.41 | 0.40 |
|  | Ensemble scGPT + GenePT-w + GenePT-s | 0.72 | 0.66 | 0.57 | 0.55 |

## Appendix D Additional visualization on the batch effect and underlying diseases biology via GenePT in the cardiomyocytes and Aorta data

In Figure D8, we visualize the original single-cell data and GenePT embeddings, coloured by disease type (top row) and patient identity (bottom row). We see that while the original data was capturing the underlying disease biology (top left; NF: non-failing heart; HCM: hearts with hypertrophic cardiomyopathy; DCM: hearts with dilated cardiomyopathy), it also was highly affected by patient batch effect (panel (b) in Figure D8; different colour indicates individual patients). On the other hand, cell embeddings generated by GenePT-s clustered primarily by disease phenotype rather than patients.

**Fig. D8:**
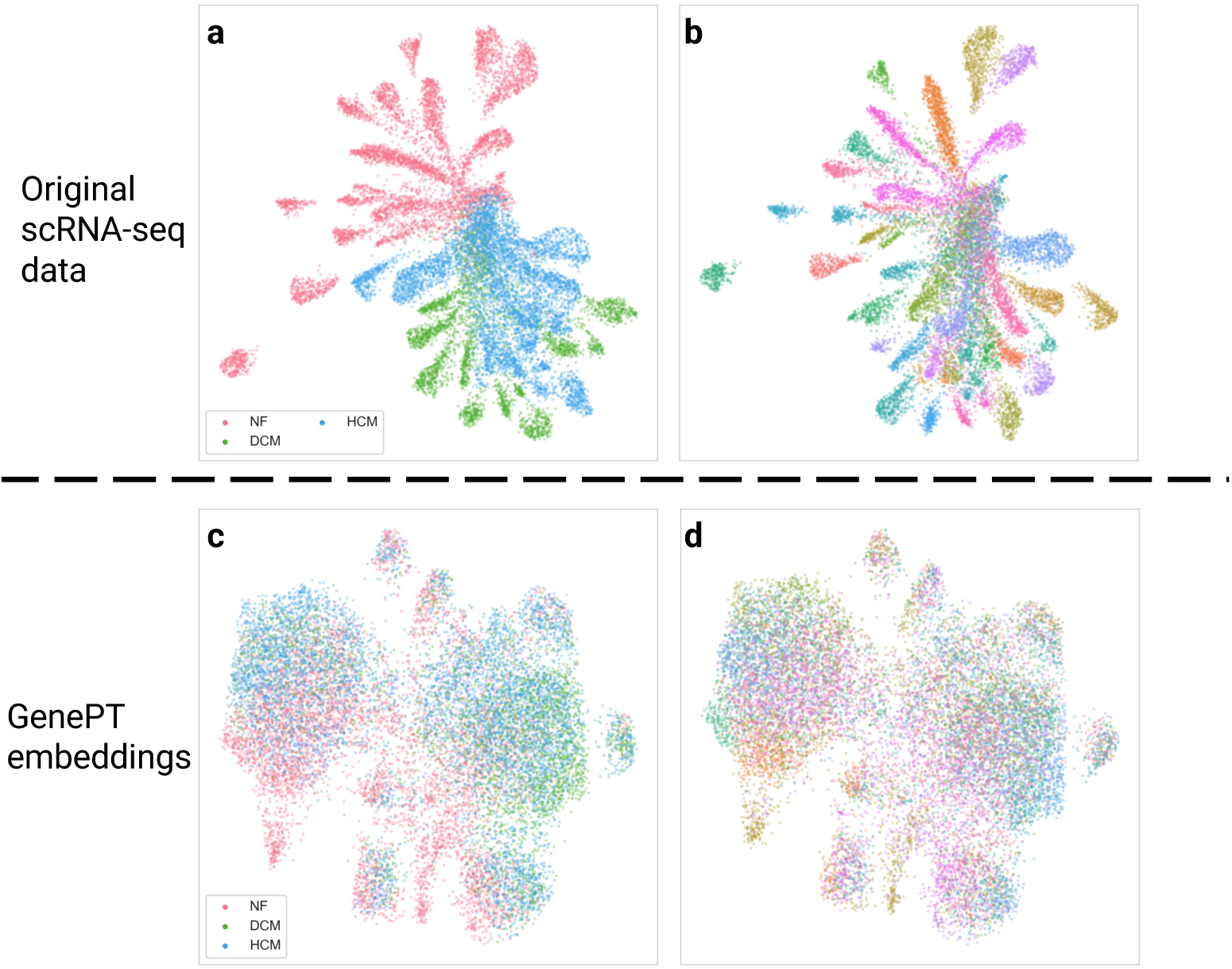
**(a)** UMAP visualization of single-cell data, coloured by disease phenotype where NF, HCM, and DCM stand for non-failing heart, hearts with hypertrophic cardiomyopathy, and hearts with dilated cardiomyopathy, respectively. **(b)** Same as **(a)**, but colored by patient id. **(c)** UMAP visualization of GenePT-s embeddings of the same set of cells as **(a)**, coloured by disease phenotype. **(d)** Same as **(c)**, but colored by patient identity.

## References

[1] Theodoris, C.V., Xiao, L., Chopra, A., Chaffin, M.D., Al Sayed, Z.R., Hill, M.C., Mantineo, H., Brydon, E.M., Zeng, Z., Liu, X.S., Ellinor, P.T.: Transfer learning enables predictions in network biology. Nature 618(7965), 616–624 (2023)

[2] Cui, H., Wang, C., Maan, H., Wang, B.: scGPT: Towards building a foundation model for single-cell multi-omics using generative AI. Nature Methods (2024)

[3] Vaswani, A., Shazeer, N., Parmar, N., Uszkoreit, J., Jones, L., Gomez, A.N., Kaiser, L- ., Polosukhin, I.: Attention is all you need. In: Proceedings of the 31st International Conference on Neural Information Processing Systems. NIPS’17, pp. 6000–6010. Curran Associates Inc., Red Hook, NY, USA (2017)

[4] OpenAI: GPT-4 technical report (2023) arXiv:2303.08774 [cs.CL]

[5] Chen, Q., Sun, H., Liu, H., Jiang, Y., Ran, T., Jin, X., Xiao, X., Lin, Z., Niu, Z., Chen, H.: A Comprehensive Benchmark Study on Biomedical Text Generation and Mining with ChatGPT (2023)

[6] Biswas, S.S.: Role of ChatGPT in public health. Annals of Biomedical Engineering 51(5), 868–869 (2023)

[7] Ayers, J.W., Poliak, A., Dredze, M., Leas, E.C., Zhu, Z., Kelley, J.B., Faix, D.J., Goodman, A.M., Longhurst, C.A., Hogarth, M., Smith, D.M.: Comparing physician and artificial intelligence chatbot responses to patient questions posted to a public social media forum. JAMA Internal Medicine 183(6), 589–596 (2023)

[8] Strong, E., DiGiammarino, A., Weng, Y., Kumar, A., Hosamani, P., Hom, J., Chen, J.H.: Chatbot vs medical student performance on Free-Response clinical reasoning examinations. JAMA Internal Medicine 183(9), 1028–1030 (2023)

[9] OpenAI: New and improved embedding model. https://openai.com/blog/new-and-improved-embedding-model. Accessed: 2023-10-4 (2023)

[10] Bommasani, R., Hudson, D.A., Adeli, E., Altman, R., Arora, S., Arx, S., Bernstein, M.S., Bohg, J., Bosselut, A., Brunskill, E., Brynjolfsson, E., Buch, S., Card, D., Castellon, R., Chatterji, N., Chen, A., Creel, K., Davis, J.Q., Demszky, D., Donahue, C., Doumbouya, M., Durmus, E., Ermon, S., Etchemendy, J., Ethayarajh, K., Fei-Fei, L., Finn, C., Gale, T., Gillespie, L., Goel, K., Goodman, N., Grossman, S., Guha, N., Hashimoto, T., Henderson, P., Hewitt, J., Ho, D.E., Hong, J., Hsu, K., Huang, J., Icard, T., Jain, S., Jurafsky, D., Kalluri, P., Karamcheti, S., Keeling, G., Khani, F., Khattab, O., Koh, P.W., Krass, M., Krishna, R., Kuditipudi, R., Kumar, A., Ladhak, F., Lee, M., Lee, T., Leskovec, J., Levent, I., Li, X.L., Li, X., Ma, T., Malik, A., Manning, C.D., Mirchandani, S., Mitchell, E., Munyikwa, Z., Nair, S., Narayan, A., Narayanan, D., Newman, B., Nie, A., Niebles, J.C., Nilforoshan, H., Nyarko, J., Ogut, G., Orr, L., Papadimitriou, I., Park, J.S., Piech, C., Portelance, E., Potts, C., Raghunathan, A., Reich, R., Ren, H., Rong, F., Roohani, Y., Ruiz, C., Ryan, J., Ré, C., Sadigh, D., Sagawa, S., Santhanam, K., Shih, A., Srinivasan, K., Tamkin, A., Taori, R., Thomas, A.W., Tramér, F., Wang, R.E., Wang, W., Wu, B., Wu, J., Wu, Y., Xie, S.M., Yasunaga, M., You, J., Zaharia, M., Zhang, M., Zhang, T., Zhang, X., Zhang, Y., Zheng, L., Zhou, K., Liang, P.: On the opportunities and risks of foundation models (2021) arXiv:2108.07258 [cs.LG]

[11] Connell, W., Khan, U., Keiser, M.J.: A single-cell gene expression language model (2022) arXiv:2210.14330 [q-bio.QM]

[12] Yang, F., Wang, W., Wang, F., Fang, Y., Tang, D., Huang, J., Lu, H., Yao, J.: scBERT as a large-scale pretrained deep language model for cell type annotation of single-cell RNA-seq data. Nature Machine Intelligence 4(10), 852–866 (2022)

[13] Lopez, R., Regier, J., Cole, M.B., Jordan, M.I., Yosef, N.: Deep generative modeling for single-cell transcriptomics. Nature Methods 15(12), 1053–1058 (2018)

[14] Lotfollahi, M., Wolf, F.A., Theis, F.J.: scgen predicts single-cell perturbation responses. Nature Methods 16(8), 715–721 (2019)

[15] Clough, E., Barrett, T.: The gene expression omnibus database. Methods in Molecular Biology 1418, 93–110 (2016)

[16] Regev, A., Teichmann, S.A., Lander, E.S., Amit, I., Benoist, C., Birney, E., Bodenmiller, B., Campbell, P., Carninci, P., Clatworthy, M., Clevers, H., Deplancke, B., Dunham, I., Eberwine, J., Eils, R., Enard, W., Farmer, A., Fugger, L., Göttgens, B., Hacohen, N., Haniffa, M., Hemberg, M., Kim, S., Klenerman, P., Kriegstein, A., Lein, E., Linnarsson, S., Lundberg, E., Lundeberg, J., Majumder, P., Marioni, J.C., Merad, M., Mhlanga, M., Nawijn, M., Netea, M., Nolan, G., Pe’er, D., Phillipakis, A., Ponting, C.P., Quake, S., Reik, W., Rozenblatt-Rosen, O., Sanes, J., Satija, R., Schumacher, T.N., Shalek, A., Shapiro, E., Sharma, P., Shin, J.W., Stegle, O., Stratton, M., Stubbington, M.J.T., Theis, F.J., Uhlen, M., Oudenaarden, A., Wagner, A., Watt, F., Weissman, J., Wold, B., Xavier, R., Yosef, N., Human Cell Atlas Meeting Participants: The human cell atlas. eLife 6 (2017)

[17] Cellxgene Data Portal. https://cellxgene.cziscience.com/docs/08Cite%20cellxgene%20in%20your%20publications. Accessed: 2023-10-4

[18] Du, J., Jia, P., Dai, Y., Tao, C., Zhao, Z., Zhi, D.: Gene2vec: distributed representation of genes based on co-expression. BMC Genomics 20, 82 (2019)

[19] Duong, D., Ahmad, W.U., Eskin, E., Chang, K.-W., Li, J.J.: Word and sentence embedding tools to measure semantic similarity of gene ontology terms by their definitions. Journal of Computational Biology 26(1), 38–52 (2019)

[20] Chen, Q., Lee, K., Yan, S., Kim, S., Wei, C.-H., Lu, Z.: BioConceptVec: Creating and evaluating literature-based biomedical concept embeddings on a large scale. PLoS Computational Biology 16(4), 1007617 (2020)

[21] Hou, W., Ji, Z.: Reference-free and cost-effective automated cell type annotation with GPT-4 in single-cell RNA-seq analysis. bioRxivy (2023)

[22] Wysocki, O., Zhou, Z., O’Regan, P., Ferreira, D., Wysocka, M., Landers, D., Freitas, A.: Transformers and the representation of biomedical background knowledge. Computational Linguistics 49(1), 73–115 (2023)

[23] Ye, R., Zhang, C., Wang, R., Xu, S., Zhang, Y.: Natural language is all a graph needs (2023) arXiv:2308.07134 [cs.CL]

[24] Sayers, E.W., Agarwala, R., Bolton, E.E., Brister, J.R., Canese, K., Clark, K., Connor, R., Fiorini, N., Funk, K., Hefferon, T., Holmes, J.B., Kim, S., Kimchi, A., Kitts, P.A., Lathrop, S., Lu, Z., Madden, T.L., Marchler-Bauer, A., Phan, L., Schneider, V.A., Schoch, C.L., Pruitt, K.D., Ostell, J.: Database resources of the national center for biotechnology information. Nucleic Acids Research 47(D1), 23–28 (2019)

[25] Levine, D., Rizvi, S.A., Lévy, S., Pallikkavaliyaveetil, N., Wu, R., Zheng, Z., Fonseca, A.O., Chen, X., Ghadermarzi, S., Dhodapkar, R.M., Dijk, D.: Cell2Sentence: Teaching Large Language Models the Language of Biology (2023)

[26] Hewitt, J., Liang, P.: Designing and interpreting probes with control tasks (2019) arXiv:1909.03368 [cs.CL]

[27] Conneau, A., Kruszewski, G., Lample, G., Barrault, L., Baroni, M.: What you can cram into a single vector: Probing sentence embeddings for linguistic properties. arXiv preprint arXiv:1805.01070 (2018)

[28] Belinkov, Y., Durrani, N., Dalvi, F., Sajjad, H., Glass, J.: What do neural machine translation models learn about morphology? arXiv preprint arXiv:1704.03471 (2017)

[29] Brown, G.R., Hem, V., Katz, K.S., Ovetsky, M., Wallin, C., Ermolaeva, O., Tolstoy, I., Tatusova, T., Pruitt, K.D., Maglott, D.R., Murphy, T.D.: Gene: a gene-centered information resource at NCBI. Nucleic Acids Research 43(Database issue), 36–42 (2015)

[30] Welcome to MyGene.py’s documentation! — MyGene.py v3.1.0 documentation. https://docs.mygene.info/projects/mygene-py/en/latest/. Accessed: 2023-10-4

[31] Seal, R.L., Braschi, B., Gray, K., Jones, T.E.M., Tweedie, S., Haim-Vilmovsky, L., Bruford, E.A.: Genenames.org: the HGNC resources in 2023. Nucleic Acids Research 51(D1), 1003–1009 (2023)

[32] Yasunaga, M., Leskovec, J., Liang, P.: Linkbert: Pretraining language models with document links. arXiv preprint arXiv:2203.15827 (2022)

[33] Luck, K., Kim, D.-K., Lambourne, L., Spirohn, K., Begg, B.E., Bian, W., Brignall, R., Cafarelli, T., Campos-Laborie, F.J., Charloteaux, B., Choi, D., Coté, A.G., Daley, M., Deimling, S., Desbuleux, A., Dricot, A., Gebbia, M., Hardy, M.F., Kishore, N., Knapp, J.J., Kovács, I.A., Lemmens, I., Mee, M.W., Mellor, J.C., Pollis, C., Pons, C., Richardson, A.D., Schlabach, S., Teeking, B., Yadav, A., Babor, M., Balcha, D., Basha, O., Bowman-Colin, C., Chin, S.-F., Choi, S.G., Colabella, C., Coppin, G., D’Amata, C., De Ridder, D., De Rouck, S., Duran-Frigola, M., Ennajdaoui, H., Goebels, F., Goehring, L., Gopal, A., Haddad, G., Hatchi, E., Helmy, M., Jacob, Y., Kassa, Y., Landini, S., Li, R., Lieshout, N., MacWilliams, A., Markey, D., Paulson, J.N., Rangarajan, S., Rasla, J., Rayhan, A., Rolland, T., San-Miguel, A., Shen, Y., Sheykhkarimli, D., Sheynkman, G.M., Simonovsky, E., Taşan, M., Tejeda, A., Tropepe, V., Twizere, J.-C., Wang, Y., Weatheritt, R.J., Weile, J., Xia, Y., Yang, X., Yeger-Lotem, E., Zhong, Q., Aloy, P., Bader, G.D., De Las Rivas, J., Gaudet, S., Hao, T., Rak, J., Tavernier, J., Hill, D.E., Vidal, M., Roth, F.P., Calderwood, M.A.: A reference map of the human binary protein interactome. Nature 580(7803), 402–408 (2020)

[34] Rolland, T., Taşan, M., Charloteaux, B., Pevzner, S.J., Zhong, Q., Sahni, N., Yi, S., Lemmens, I., Fontanillo, C., Mosca, R., Kamburov, A., Ghiassian, S.D., Yang, X., Ghamsari, L., Balcha, D., Begg, B.E., Braun, P., Brehme, M., Broly, M.P., Carvunis, A.-R., Convery-Zupan, D., Corominas, R., Coulombe-Huntington, J., Dann, E., Dreze, M., Dricot, A., Fan, C., Franzosa, E., Gebreab, F., Gutierrez, B.J., Hardy, M.F., Jin, M., Kang, S., Kiros, R., Lin, G.N., Luck, K., MacWilliams, A., Menche, J., Murray, R.R., Palagi, A., Poulin, M.M., Rambout, X., Rasla, J., Reichert, P., Romero, V., Ruyssinck, E., Sahalie, J.M., Scholz, A., Shah, A.A., Sharma, A., Shen, Y., Spirohn, K., Tam, S., Tejeda, A.O., Wanamaker, S.A., Twizere, J.-C., Vega, K., Walsh, J., Cusick, M.E., Xia, Y., Barabási, A.-L., Iakoucheva, L.M., Aloy, P., De Las Rivas, J., Tavernier, J., Calderwood, M.A., Hill, D.E., Hao, T., Roth, F.P., Vidal, M.: A proteome-scale map of the human interactome network. Cell 159(5), 1212–1226 (2014)

[35] Greene, C.S., Krishnan, A., Wong, A.K., Ricciotti, E., Zelaya, R.A., Himmelstein, D.S., Zhang, R., Hartmann, B.M., Zaslavsky, E., Sealfon, S.C., et al.: Understanding multicellular function and disease with human tissue-specific networks. Nature Genetics 47(6), 569–576 (2015)

[36] Consortium, U.: Uniprot: the universal protein knowledgebase in 2023. Nucleic Acids Research 51(D1), 523–531 (2023)

[37] Luecken, M.D., Büttner, M., Chaichoompu, K., Danese, A., Interlandi, M., Mueller, M.F., Strobl, D.C., Zappia, L., Dugas, M., Colomé-Tatché, M., Theis, F.J.: Benchmarking atlas-level data integration in single-cell genomics. Nature Methods 19(1), 41–50 (2022)

[38] Blondel, V.D., Guillaume, J.-L., Lambiotte, R., Lefebvre, E.: Fast unfolding of communities in large networks. Journal of Statistical Mechanics: Theory and Experiment 2008(10), 10008 (2008)

[39] Li, Y., Ren, P., Dawson, A., Vasquez, H.G., Ageedi, W., Zhang, C., Luo, W., Chen, R., Li, Y., Kim, S., et al.: Single-cell transcriptome analysis reveals dynamic cell populations and differential gene expression patterns in control and aneurysmal human aortic tissue. Circulation 142(14), 1374–1388 (2020)

[40] Alsaigh, T., Evans, D., Frankel, D., Torkamani, A.: Decoding the transcriptome of calcified atherosclerotic plaque at single-cell resolution. Communications Biology 5(1), 1084 (2022)

[41] Chou, C.-H., Jain, V., Gibson, J., Attarian, D.E., Haraden, C.A., Yohn, C.B., Laberge, R.-M., Gregory, S., Kraus, V.B.: Synovial cell cross-talk with cartilage plays a major role in the pathogenesis of osteoarthritis. Scientific Reports 10(1), 10868 (2020)

[42] Cheng, S., Li, Z., Gao, R., Xing, B., Gao, Y., Yang, Y., Qin, S., Zhang, L., Ouyang, H., Du, P., Jiang, L., Zhang, B., Yang, Y., Wang, X., Ren, X., Bei, J.-X., Hu, X., Bu, Z., Ji, J., Zhang, Z.: A pan-cancer single-cell transcriptional atlas of tumor infiltrating myeloid cells. Cell 184(3), 792–80923 (2021)

[43] Schirmer, L., Velmeshev, D., Holmqvist, S., Kaufmann, M., Werneburg, S., Jung, D., Vistnes, S., Stockley, J.H., Young, A., Steindel, M., Tung, B., Goyal, N., Bhaduri, A., Mayer, S., Engler, J.B., Bayraktar, O.A., Franklin, R.J.M., Haeussler, M., Reynolds, R., Schafer, D.P., Friese, M.A., Shiow, L.R., Kriegstein, A.R., Rowitch, D.H.: Neuronal vulnerability and multilineage diversity in Multiple Sclerosis. Nature 573(7772), 75–82 (2019)

[44] Chaffin, M., Papangeli, I., Simonson, B., Akkad, A.-D., Hill, M.C., Arduini, A., Fleming, S.J., Melanson, M., Hayat, S., Kost-Alimova, M., et al.: Single-nucleus profiling of human dilated and hypertrophic cardiomyopathy. Nature 608(7921), 174–180 (2022)

[45] Pedregosa, F., Varoquaux, G., Gramfort, A., Michel, V., Thirion, B., Grisel, O., Blondel, M., Prettenhofer, P., Weiss, R., Dubourg, V., Vanderplas, J., Passos, A., Cournapeau, D., Brucher, M., Perrot, M., Duchesnay, E.: Scikit-learn: Machine learning in Python. Journal of Machine Learning Research 12, 2825–2830 (2011)

[46] Bruford, E.A., Braschi, B., Denny, P., Jones, T.E.M., Seal, R.L., Tweedie, S.: Guidelines for human gene nomenclature. Nature Genetics 52(8), 754–758 (2020)

[47] AI4Science, M.R., Quantum, M.A.: The impact of large language models on scientific discovery: a preliminary study using gpt-4. arXiv preprint arXiv:2311.07361 (2023)

[48] Kusupati, A., Bhatt, G., Rege, A., Wallingford, M., Sinha, A., Ramanujan, V., Howard-Snyder, W., Chen, K., Kakade, S., Jain, P., et al.: Matryoshka representation learning. Advances in Neural Information Processing Systems 35, 30233–30249 (2022)

[49] Rives, A., Meier, J., Sercu, T., Goyal, S., Lin, Z., Liu, J., Guo, D., Ott, M., Zitnick, C.L., Ma, J., et al.: Biological structure and function emerge from scaling unsupervised learning to 250 million protein sequences. Proceedings of the National Academy of Sciences 118(15), 2016239118 (2021)

[50] Visscher, P.M., Brown, M.A., McCarthy, M.I., Yang, J.: Five years of gwas discovery. The American Journal of Human Genetics 90(1), 7–24 (2012)

